# Functional characterization of five triterpene synthases through De-novo assembly and transcriptome analysis of *Euphorbia grantii* and *Euphorbia tirucalli*

**DOI:** 10.1101/2023.04.05.535548

**Authors:** Ashish Kumar, Dhanashri S. Mulge, Kalyani J. Thakar, Avinash Pandreka, Amruta D. Warhekar, Sudha Ramkumar, Poojadevi Sharma, Sindhuri Upadrasta, Dhanasekaran Shanmugam, Hirekodathakallu V. Thulasiram

## Abstract

*Euphorbia grantii* and *Euphorbia tirucalli* known to synthesize diverse triterpenes including euphol and tirucallol. These two triterpenes known to possess potent anti-cancer, anti-bacterial, and anti-fungal properties along with various other biological activities. In this study, De-novo assembly and comparative transcriptome analysis of leaf and stem tissues of *E. tirucalli* and *E. grantii* were carried out to identify thirteen triterpene synthases from 1,40,227 in correlation with the metabolic profiling. Comparative transcriptome analysis indicated that EutTTS4 and EutTTS5 genes which encodes for euphol/tirucallol and tirucallol synthase were highly expressed in leaf and stem tissue. The genes which encodes α-amyrin synthase (EutTTS1) and lupeol synthase (EutTTS2) were characterized by overexpressing them in YPH499 yeast strain. We have developed using hem1 and erg7 knock yeast strain of lanosterol deficient yeast (TMBL17) and used for over expression of friedelin synthase (EutTTS3), and two novel triterpenes synthases such as euphol/tirucallol synthase (EutTTS4) and tirucallol synthase (EutTTS5). These results are very useful in large scale production of triterpenes by genomic integration of respective triterpene synthases in TMBL yeast strain developed in this study.

**Significance Statement:** We have functionally characterized triterpene synthases from *E. tirucalli* and *E. grantii* and developed a hem1 and erg7 knock out of lanosterol deficient yeast (TMBL17) for the large-scale production of triterpene and triterpene related products.

## 1.0 Introduction

More than 20,000 triterpenoids with over 150 unique carbon skeletons are known to date. They are known to possess wide range of medicinal properties and being used in traditional medicine as well as in Ayurveda, and they have high commercial value. Applanatum triterpenoids and polyketides shown to possess anti-cancer effect whereas Lanostanoid triterpenes were found to be potent antioxidant and neuro-protective properties (Liu et al., 2018). The triterpenoids play many physiological roles such as plants contain cycloartenol which is mainly involved in the biosynthesis of sterols, whereas β-amyrin is involved in the defense mechanism. When the plant is having physical injury, the plant releases the methyl jasmonate hormone resulting in increased production of β-amyrin-based saponins (Vernoud et al., 2021). Likewise in case of animals, lanosterol is responsible for the biosynthesis of sterols (Ohyama et al., 2009). Triterpenoids are one of the major classes of isoprenoids constructed from six isoprene units. These are generated from isopentenyl pyrophosphate (IPP) and dimethylallyl pyrophosphate (DMAPP) biosynthesized from mevalonic acid pathway (MVA) and methyl erythritol pyrophosphate pathway (MEP). Further, DMAPP and IPP combine head to tail manner to produce monoterpenes such as geranyl pyrophosphate (GPP) catalyzed by geranyl pyrophosphate synthase. GPP acts as a precursor for the biosynthesis of monoterpenes such as linalool, limonene etc. Farnesyl diphosphate synthase catalyzes the condensation one molecule IPP with GPP to produce farnesyl pyrophosphate (FPP), which is the branching intermediate involved in the biosynthesis of sesquiterpenoids, diterpenoids, farnesylated proteins, and triterpenoids. Sesquiterpene synthases convert FPP in to various sesquiterpenes through intramolecular carbocation rearrangements of farnesyl carbocation. On the other hand, squalene synthase catalyzes the head-to-head condensation of two molecules of FPP to form squalene through pre-squalene as intermediate (Thulasiram and Poulter, 2006, Thulasiram et al., 2007),. Squalene is a precursor for the biosynthesis of various acyclic triterpenes such as ekeberins, botryococcine etc whereas cyclic triterpenes are biosynthesized from (*S*)-2,3-oxidosqualene which forms through oxidation of squalene- by-squalene epoxidase (Niehaus et al., 2012, Niehaus et al., 2011). In prokaryotes, hopene is the only cyclic triterpene synthesized directly from squalene. On the other hand, in most of the eukaryotes, (*S*)-2,3-oxidosqualene undergoes reduction of the epoxy group at C2 and C3 position by Asp522 residue of triterpene synthases, thus initiating the cyclization step and further arrangements to synthesize cyclic triterpenes (Corey et al., 1997). Cyclization of (*S*) 2,3-oxidosqualene leads to the formation of various triterpenes through two carbocation intermediates *viz.* Protosteryl cation (PC) and Dammarenyl cation (DC). Tetracyclic triterpenes such as cycloartenol and lanosterol formed through PC whereas, DC is involved as an intermediate in the formation of tetracyclic triterpenes such as euphol and tirucallol as well as pentacyclic triterpenes such as, α-amyrin, β- amyrin, lupeol etc (Liu et al., 2018) (Figure S29).

In this study, we have quantified the triterpenes from the leaf and stem tissues of *E. tirucalli* and *E. grantii*. This data was corelated with the transcriptome analysis to identify thirteen triterpene synthases. Out of which, five were functionally characterized as α-amyrin synthase, lupeol synthase, friedline synthase, and two novel triterpene synthases such as euphol/tirucallol synthase and tirucallol synthase. We also developed hem1 and erg7 knock out of lanosterol deficient yeast (TMBL17) for the functional characterization of triterpene synthases. This strain can also be utilized for the large-scale production of triterpene and triterpene related products by using the synthetic biology tools.

## 2.0 Result and discussion

### 2.1 Triterpenoid profiling of different tissues

The study carried out differential tissue metabolite profiling and quantification of targeted metabolites in *E. tirucalli* and *E. grantii* leaf and stem tissues. GC-MS analysis revealed that triterpenoid and steroid metabolites together represented 78-83 % of the total metabolite content of *E. tirucalli* and *E. grantii* leaf and stem tissues of which triterpenoid content was 60-70 % and steroidal content was 10-20 %, respectively, as shown in (Figure S2A). Further, *E. tirucalli* leaf had the highest triterpenoid content, while *E. grantii* stem tissue had the highest steroid content. Triterpenoids, tirucallol, euphol, β-amyrin, α-amyrin, lupeol, cycloartenol, and friedelin were confirmed in the leaf and stem tissues of *E. tirucalli* and *E. grantii* by GC-MS analyses by comparing the retention time, mass fragmentation pattern and co-injection studies using respective reference compounds. These metabolites were quantified, and high content variation was evident among *Euphorbia* plants in stem and leaf tissues (Figure 1D). Among them, tirucallol and euphol emerged as major metabolites accounting for 45-66% of the total triterpenoid content of *E. tirucalli* and *E. grantii* plants. Further, tirucallol and euphol were present in the 4: 1 ratio in *E. tirucalli* leaf and stem tissues and the 1: 1 ratio in *E. grantii* leaf and stem tissues, respectively. In general, leaf tissue had higher tirucallol and euphol content than stem tissue for both plants. Among other triterpenoids, α-amyrin, cycloartenol and β-amyrin were highly present in the leaf of *E. grantii* whereas, cycloartenol was detected in *E. tirucalli* leaf and stem tissues in very trace amount. Further, Lupeol was found highly in *E. tirucalli* leaf then in stem tissues, whereas it was detected only in fraction in both leaf and stem tissue of *E. grantii.* Friedelin was highly present in *E. tirucalli* leaf then in other tissues. These results suggest that triterpenoid content vary with tissue type in *E. tirucalli* and E. *grantii*. Concerning steroids, β-sitosterol was the most abundant sterol produced in *E. tirucalli* and *E*. *grantii* tissues accounting for 50-80% of total steroid content. β-sitosterol content was high in the leaf then stem tissues and it accounted for 80% of *E. grantii* stem extract (Figure 1D, S2B). Metabolite profiling and quantification of triterpenoids in targeted plant tissues was compared with differential expression of transcriptome from leaf and stem tissues of *E. tirucalli* and *E. grantii* to identify the novel triterpene synthases.

**Fig 1.**
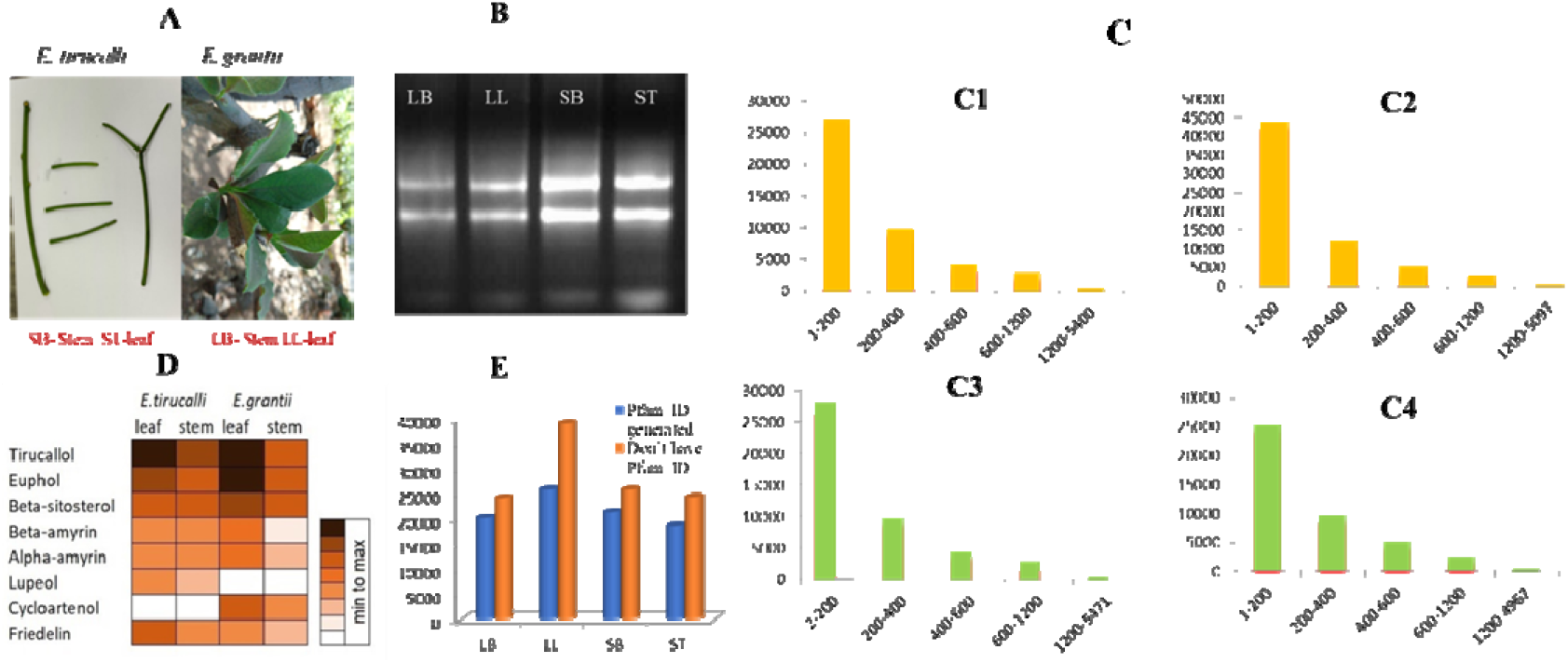
Triterpene metabolite profiling and transcriptome analysis of leaf and stem tissues of *E. grantii* and *E. tirucalli*. (A) *E. tirucalli* and *E. grantii* leaf and stem. (B) Agarose gel image of RNA isolated from the different tissues of euphorbia. C (C1, C2, C3 and C4) this is a virtual ribosome analysis of LB, LL, ST, and SB. (D) metabolite profiling of different triterpenes in leaf and stem tissues. (E) the number of Pfam IDs generated from the transcripts of four tissues.

### 2.2 Transcriptome analysis

*De-novo* assembly and transcriptome analysis of 4 samples ST, SB, LB, and LL of Euphorbia genus was carried out. On an average of 29.73 million, high-quality raw reads were generated. These raw reads were quality checked using FastQc (Figure S3, S4). Raw reads were trimmed of base pairs present on the terminal of the sequences with a low Phred score to generate processed reads using a trimmomatic tool (Table 1). Trinity was used to generate a full-length transcript by the de-novo assembly of raw reads. Cd-hit was used to remove duplicate transcripts to create a master sequence (Table 2). Clustering transcripts from the 4 samples resulted in 140227 unique representative transcripts (average length of 1148bp and N50 of 1631bp) (Figure S26). ∼72.51% of transcripts were functionally annotated using the public database, and KO_IDs were assigned on the KEGG server (Table 2) (Figure S1 A, B). Simple sequence repeat analysis revealed that ∼23% of the transcripts were found to have SSRs. Differential expression analysis revealed that ∼16.29% of transcripts were differentially regulated in different combinations of DGE comparisons. Total clustered transcripts from *E. tirucalli* and *E. grantii* tissues leaf and stem were submitted to virtual ribosome generated protein sequences ORF (Figure 1C1-C4) (Table 2). These protein sequences were further submitted for the Pfam analysis, which has assigned to 13608-ST, 21260-SB, 25802-LL, and 20132- LB Pfam_ID to the protein sequences of *E. tirucalli* and *E. grantii* leaf and stem tissues (Figure 1E). Pfam_ID PF13243 (SQHop_cyclase_C) and PF13249 (SQHop_cyclase_N) designate the conserved domain for triterpene synthases, these two ids were searched in the four tissues, and 9 triterpene synthases from LB, 4 triterpene synthases from LL, 9 triterpene synthases from SB, 10 triterpene synthases from ST were identified (Table S1-S4). Further, these triterpene synthases were aligned in clustal omega, and the sequences with 98% or above identity were removed to identify thirteen unique triterpene synthases (Table 3). Functional annotation of identified triterpenes was carried out through NCBI blast analysis. Five transcripts show the highest match with β- amyrin synthase, four with lupeol synthase, and four with putative oxido-squalene synthases. Nine HMGR were identified from *E. grantii* and *E. tirucalli* based on Pfam ID PF00368 (HMG-CoA_red) and three DXS having the following Pfam ID PF13292 (DXP_synthase_N) PF02779 (Transket_pyr) PF02780 (Transketolase_C). Expression analysis of these genes by real-time PCR has led to the prediction of which pathway is involved in the biosynthesis of triterpenes. One FDS is identified in all of the tissues of *E. grantii* and *E. tirucalli*.

**Table 1.** Showing sequencing reads generated after Illumina sequencing.

| Tissue | Raw reads (in millions) | Processed reads (in millions) |
| --- | --- | --- |
| LB | 28.64 | 26.02 |
| LL | 30.83 | 27.86 |
| SB | 29.74 | 26.87 |
| ST | 31.2 | 28.25 |

**Table 2.** The general statistics of assembled transcript analysis for the generation of Pfam_ID.

|  | LB | LL | SB | ST |
| --- | --- | --- | --- | --- |
| Total no of transcript | 44260 | 64694 | 45665 | 42938 |
| Transcripts were given KO_ID | 5781 | 6796 | 5892 | 5971 |
| Transcripts having ORF | 44259 | 64691 | 45665 | 42937 |
| Transcripts have Pfam_ID | 20132 | 25802 | 21260 | 13608 |

**Table 3.** Identified triterpene synthase from *E. grantii* and *E. tirucalli* transcriptome.

| Transcript I. D | Description of the transcript | Identity | Query coverage | Transcript length |
| --- | --- | --- | --- | --- |
| ST_c28604_g2_i2 (EutTTS1) | β-amyrin synthase [*Lotus japonicus*] | 82.1 | 95.25 | Full length |
| ST_c27781_g2_i2 (EutTTS4) | lupeol synthase [*Ricinus communis*] | 76.9 | 87.96 | Full length |
| ST_c27781_g2_i6 (EutTTS3) | putative oxidosqualene cyclase [*Euphorbia tirucalli*] | 79.4 | 45.72 | Full length |
| ST_c13055_g1_i1 (EutTTS8) | putative oxidosqualene cyclase [*Euphorbia tirucalli*] | 80.3 | 100 | 510 bp from N-terminal and 1.2 kb from C-terminal |
| SB_c36763_g3_i6 | lupeol synthase [*Ricinus communis*] | 83.6 | 96.05 | 180 bp from N-terminal and 150 bp from C-terminal |
| SB_c36763_g3_i3 (EutTTS2) | lupeol synthase [*Ricinus communis*] | 83.6 | 96.05 | Full length |
| LL_c40919_g7_i5 | β-amyrin synthase [*Eleutherococcus senticosus*] | 58.8 | 84.58 | Full length |
| LL_c41931_g2_i7 (EugTTS7) | β-amyrin synthase [*Euphorbia tirucalli*] | 91.2 | 72.61 | 360 bp from C-terminal |
| LL_c45609_g2_i2 | putative oxidosqualene cyclase [*Euphorbia tirucalli*] | 78.6 | 97.24 | 600 bp missing from C-terminal |
| LB_c34999_g1_i1 (EugTTS6) | β-amyrin synthase [*Euphorbia tirucalli*] | 96 | 84 | Full length |
| LB_c35995_g3_i3 (EugTTS5) | β-amyrin synthase [*Bupleurum chinense*] | 57 | 85 | Full length |
| LB_c36778_g2_i1 | putative oxidosqualene cyclase [*Euphorbia tirucalli*] | 88 | 99 | C-terminal 300bp and N-33bp |
| LB_c36778_g2_i3 | lupeol synthase [*Ricinus communis*] | 77 | 88 | Full length |

### 2.3 Phylogenetic and conserved amino acid analysis

The ORF of thirteen unique triterpene synthases from *E. tirucalli* (Eut) and *E. grantii* (Eug) was aligned against the ORF of triterpene synthase retrieved from UniProt using MEGA5. The phylogenetic tree was generated by the neighbor-joining method. This data was used to predict the function of triterpene synthase identified after transcriptome analysis. Transcript LL_c41931_g2_i2 and LB_c34999_g1_i1 match more than 90 % with the β- amyrin synthase from *E. tirucalli* (Figure S27) (Kajikawa et al., 2005). These two transcripts also have MWCYCR conserved amino acid on their active site, which shows that these two transcripts are involved in the biosynthesis of pentacyclic triterpenes (Figure S27). ST_c13055_g1_i1 matches cycloartenol synthase, whose identity is more than 95% with *Ricinus communis* and 90% with *Arabidopsis thaliana*; amino acid residue H is present on the 260 positions in the ORF instead of Y, which will neutralize the carbocation generated on 20 carbons stopping the formation of the fifth ring structure. It is present in all the cycloartenol synthases functionally characterized to date (Figure S27). LB_c35995_g3_i3 and LL_c409191_g7_i5 match more than 55% with the Dammrenediol synthase II from *Panax ginseng*. The ORF of these two unique triterpenes were analyzed and found a different amino acid at 257 and 260 positions, L and F are present in LB_c35995_g3_i3 and L at 259 and 260 part is present in LL_c40919_g7_i5 instead of M and Y, the conserved amino acids present in most of the triterpenes (Figure S27). These active sites amino acid variation in LL_c40919_g7_i5 and LB_c35995_g3_i3 may lead to tetracyclic triterpenes (Euphol or tirucallol synthases). ST_c28604_g2_i2 transcript ORF has the highest matches with tirucalla-7,24-diene-3β-ol synthase gene from *A. indica,* which was functionally characterized in our lab, and its product has structural similarity with the Euphol and tirucallol. SB_c36763_g3_i3, SB_c36763_g3_i6 and ST_c27781_g2_i6 is very close to the lupeol synthase from *Arabidopsis thaliana*. LB_c36778_g2_i10 and LB_c36778_g2_i3 is very close to isomultiflorenal synthase. LB_c36778_g2_i1 is very close to β-amyrin synthase EUPTII from *E. tirucalli* (Figure 2). After transcriptome analysis, phylogenetic analysis, and active site amino acid analysis, we choose five triterpene synthases for the cloning and functional characterization in the yeast system.

**Fig 2.**
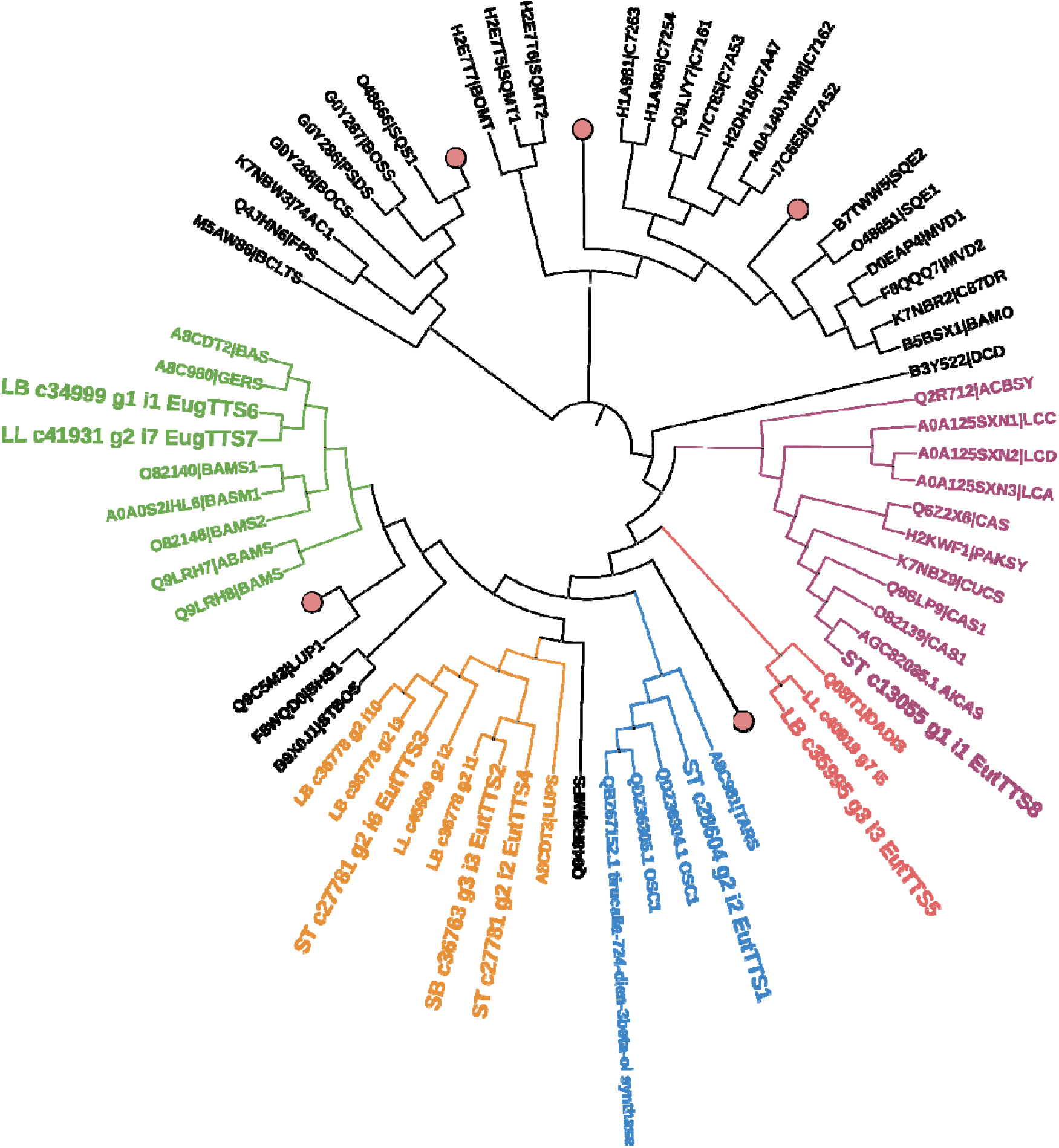
Phylogenetic analysis of the triterpene cyclases identified from the *E. tirucalli* and *E. grantii* transcriptome analysis. Purple-coloured leaves show the triterpenes synthesized through Protosteryl cation; pink-coloured leaves show the triterpenes synthesized through Dammarane cation. The blue coloured leaf shows the triterpenes biosynthesizing through Dammarane cation, whereas EutTTS1 is biosynthesizing through ursane carbocation. Orange colour leaves show the triterpenes biosynthesizing through lupenyl cation, and green colour leaf’s are leading the triterpene biosynthesizing through olenyl cation. The leaf’s in black colours are mainly the non-cyclic triterpene except LUP1 which is a pentacyclic triterpene.

### 2.4 CRISPR mediated knockout of erg7 and hem1 gene in Yeast (YPH499)

Yeast (YPH499) MATa ura3-52 lys2-801_amber ade2-101_ochre trp1-Δ63 his3-Δ200 leu2- Δ1 is a haploid strain having lanosterol synthase. This triterpene synthase utilizes (*S*) 2,3-oxidosqualene as a substrate. Lanosterol synthase on utilization of 2-3, oxidosqualene interrupts the function of heterologously expressed triterpene synthase as it uses the same substrate. To enhance the activity of heterologously expressed triterpene synthase, it is necessary to knock out lanosterol synthase. This gene is essential for the survival of the yeast. We have provided ergosterol as an external supplement to compensate for the lanosterol. We have also knocked out the hem1 gene to grow the lanosterol synthase deficient yeast strain aerobically, using Gill77 strain as a reference. CRISPR vector pML107 has been used to knock out hem1 on chromosome IV. The following sequence was targeted CCGCACCACATGCGAAAAATGGC at position 77 on the ORF of hem1. To target the given sequence gRNA, GCCATTTTTCGCAT GTGGTGCGG was designed and cloned on the MCS of the pML107 vector. This construct was transformed using S.c easy competent cell preparation kit in YPH499 yeast strain. The leu2 cassette (Figure S35 supplementary) has promoter, terminator, and specific components needed to express the leu2 gene, selection of the knockout colony was performed on CSM- w/o- leu-ura. This leu2 cassette has 40 bp overhangs, the same as the sequences on both sides of the target sequence. These overhangs are responsible for the heterologously directed recombination at the target site on 106 positions in the two positive colonies. This hem1 deficient yeast strain was screened based on colony PCR (Figure S22), metabolite profiling (Figure 6), and sequencing from the genome of yeast isolated from the specific colony (Figure S24). This leads to the knockout of the hem1 gene and the integration of the leu2 cassette at the IV chromosome. This hem1 deficient strain was grown on YPD and made competent using Sc easy competent cell preparation kit. CRISPR vector pML107 has been used to knock out erg7 on chromosome VIII. The following sequence was targeted TGAGCTAGGCCGAGAAAGCTGGG at position 72 on the ORF of erg7. To target the given position, gRNA TGAGCTAGGCCGAGAAAGCTGGG was designed and cloned on the MCS of the pML107 vector. This construct was transformed using S.c easy competent cell kit in the YPH499 yeast strain. The trp1 cassette (Figure S23) has promoter, terminator, and specific components needed to express the trp1 gene, selection of the knockout colony was performed on CSM- w/o-ura-trp. This trp1 cassette has 40 bp overhangs, the same as the sequences on both sides of the target sequence. These overhangs are responsible for the heterologously directed recombination at the target site 137 positions of the ORF for the two positive colonies 19 and 21. The knockout of hem1 and erg7 genes and the integration of leu2 and trp1 cassette at IV and VIII chromosomes of the yeast was confirmed by the Sanger sequencing. A double knockout confirmed colony was grown in YPD having external supplement 13µg/ml hemin chloride and 20µg/ml ergosterol. Total metabolite extracted from the double knockout strain was analyzed on GC-MS shows no peaks of lanosterol compared to the control (Figure 6).

### 2.5 Heterologous expression and functional characterization of EutTTS (*Euphorbia tirucalli* triterpene synthase), and EugTTS (*Euphorbia grantii* triterpene synthase)

We have identified thirteen triterpene synthases after transcriptome analysis. Out of thirteen triterpene synthases, we chose five full-length triterpene synthase for the functional characterization (Table 3). Open reading frames of the five transcripts were cloned in pYES2/CT vector at multiple cloning sites. EutTTS1 gene with the transcript length of 2.292 kb (Figure S7 lane2) and calculated molecular weight protein is 87.69 kD. This gene is in the same clade as the genes biosynthesizing tirucalla-7,24-dien-3-β-ol (Hodgson et al., 2019, Pandreka et al., 2021), a tetracyclic product in the phylogenetic tree. Whereas it has Y at 260 positions (Figure S27), which favors the formation of the pentacyclic product. This gene matches 82% with the β- amyrin synthase from *Ricinus communis* and 83% *Manihot esculenta(Kushiro et al., 2000)*. This gene was cloned in pYES2/CT and transformed in INVISc1 yeast cell (MATa/α his3D1 leu2 trp1-289 ura3-52) and induced with 2% galactose. The induced yeast pellet was saponified and extracted thrice with the n-hexane and analyzed GC-MS (Figure S8). The α amyrin peak was confirmed by mass spectrum (Figure S14), retention time, and co-injection with the authentic standard (Figure S7, S21). The EutTTS2 transcript is of 2.283 kilobase pairs (Figure S7 lane 6), and the calculated weight of the protein is 88.87 kD. This transcript is very close to the leaf, indicating lupeol synthase from *Bruguiera gymnorhiza* (Basyuni et al., 2007) in phylogeny and matches 98% with lupeol synthase from *Ricinus communis* (Guhling et al., 2006). The protein also has LFCYCR instead of the MWCYCR motif at the active site (Figure S27). The ORF was cloned in pYES2/CT, transformed into yeast INVISc1 cells, and induced with 2% galactose. The induced culture pellet was saponified and extracted thrice with the n-hexane and analyzed GC-MS. The lupeol peak was confirmed by mass spectrum (Figure S15), retention time, and co-injection with the authentic standard (Figure 4, Figure S9). The EutTTS3 is another triterpene cyclase identified from *E. tirucalli* and has a 2.283 kilobase pair (Figure S7 lane 11) open reading frame with a protein weight of 87.78 kD. This transcript matches 75.76% with the lupeol synthase from *Ricinus communis* (Guhling et al., 2006) and is cloned in the pYES2/CT vector. This was transformed into a TMBL17 yeast strain and induced with 2% galactose at 30 □C for 24 hr. The induced pellet was saponified and extracted thrice with the n-hexane and analyzed GC-MS. Based on the mass spectrum (Figure S16), retention time, and co-injection studies with authentic standards, the friedelin peak was confirmed (Figure 5, Figure S10). EutTTS4 is the triterpene cyclase of 2.2 kb (Figure S7 lane 16) identified from the *E. tirucalli* and matches 81% with the lupeol synthase from *Ricinus communis* (Guhling et al., 2006). In contrast, phylogeny has confirmed the presence of lupeol synthase from *Bruguiera gymnorhiza* (Basyuni et al., 2007) in the same clade (Fig 5). The ORF was cloned into a pYES2/CT vector, transformed into TMBL17, and induced with 2% galactose at 30 □C for 24 hours. The induced pellet was saponified and extracted thrice with the n-hexane and analyzed GC-MS. The euphol and tirucallol peaks were confirmed based on the mass spectrum (Figure S17), retention time, and co-injection with the authentic standard purchased from ChemFaces (Figure 6, Figure S11). The EugTTS5 has been identified from *E. grantii* stem and has 2.274 kilobase pair (Figure S7 lane 19) with a protein weight of 87.81 kD. This gene matches 98.06% with the euphol and tirucallol synthase from *E. tirucalli (Qiao et al., 2021)*and 82.61% with the butyrospermol from *Euphorbia lathyris* (Giner and Djerassi, 1995) after Blastp analysis and is also present in the same clade as Dammarenediol synthase II from *Panax ginseng*(Giner *and Djerassi, 1995)* in the phylogeny (Figure 5). This gene has F at the 260 positions instead of Y (Figure S27). This helped us predict this gene to biosynthesize either euphol or tirucallol. ORF of this gene was cloned into pYES2/CT vector and induced with 2% galactose at 30 □C for 24 hr. The induced pellet was saponified and extracted thrice with the n-hexane and analyzed GC-MS. The tirucallol peak was confirmed by mass spectrum (Figure S18), retention time, and co-injection with the authentic standard purchased from ChemFaces (Figure 7, Figure S12). The accession number of these TTS are EutTTS1-MH880792.1, EutTTS2-MH880793.1, EutTTS3-MK387072.1, EutTTS4-MK387071.1, and EugTTS5-MK387070.1.

**Fig 3.**
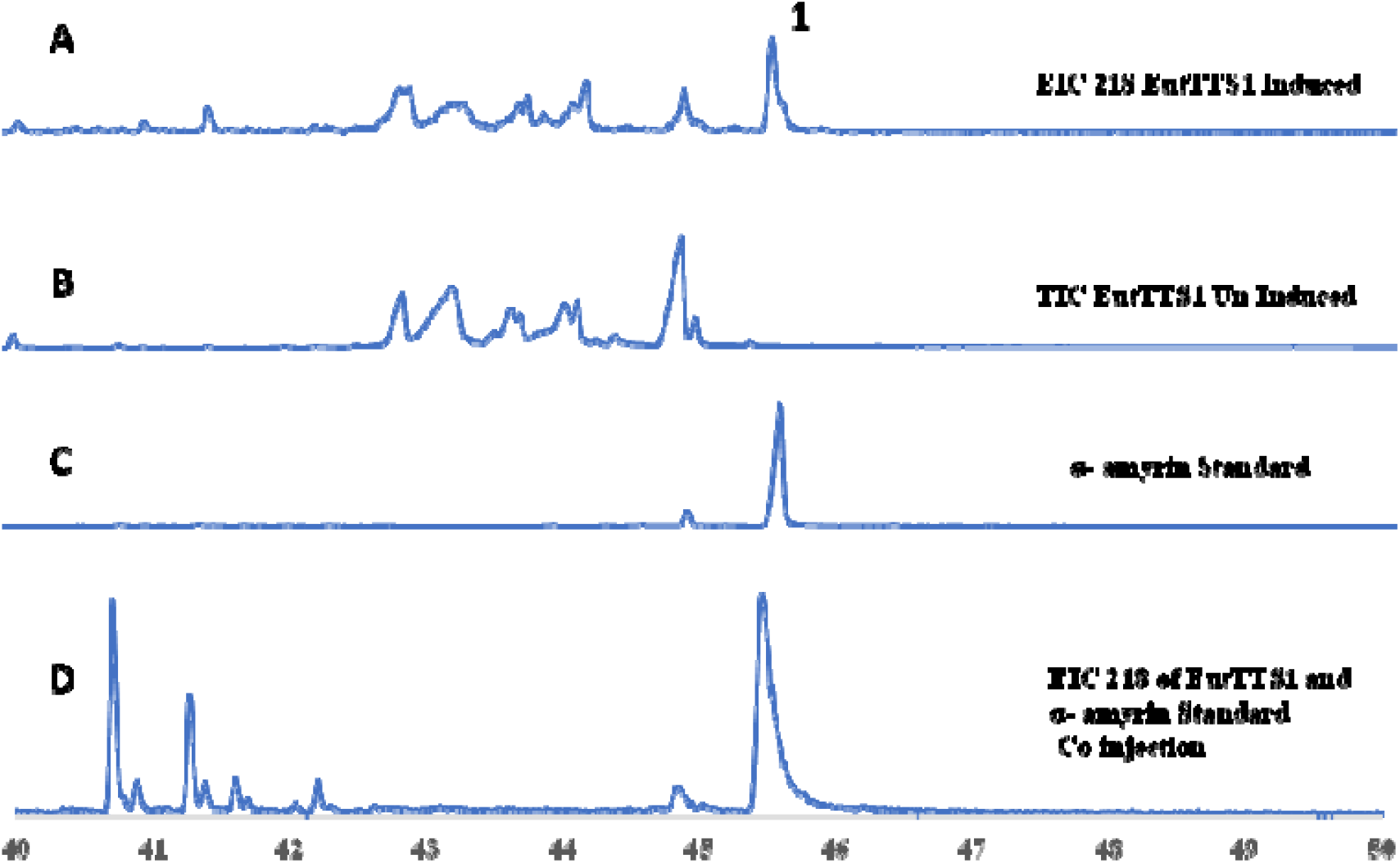
EIC (Extracted ion chromatogram) drawn from the TIC of total metabolites extracted from yeast (YPH499 strain) harbouring EutTTS1 in pYES2/CT vector. EIC was obtained having the m/z value for 218 ion. A) On comparing EIC of total metabolites after induction with 2% galactose and B) un-induced with 2% dextrose yeast samples were compared with the C) alpha amyrin standard, further D) co-injection analysis was performed to confirm the peak 1 is the product of the EutTTS1 gene.

**Fig 4.**
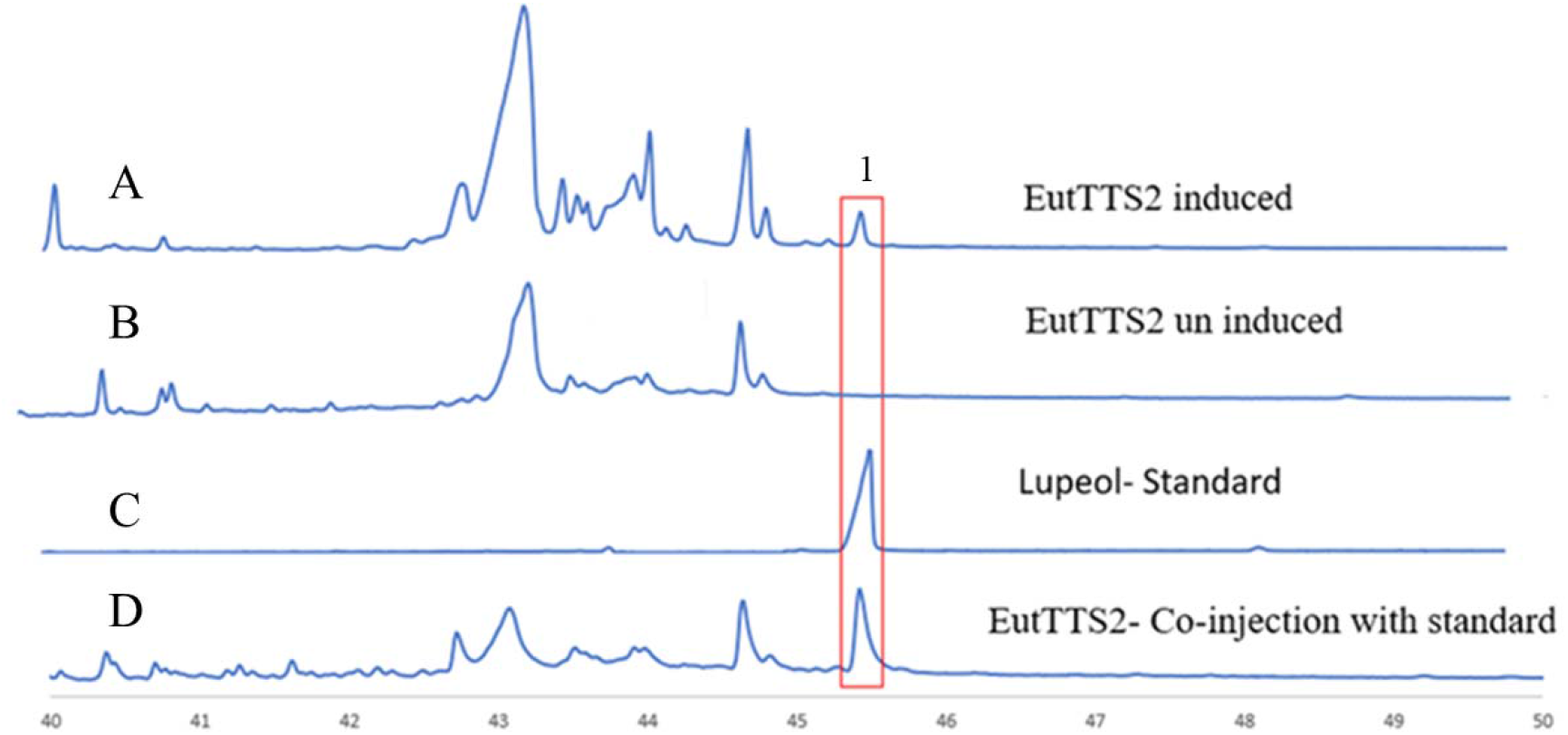
TIC (Total ion chromatogram) of total metabolites extracted from yeast harboring EutTTS2 in pYES2/CT vector. A) On comparing TIC of total metabolites after induction with 2% galactose and B) un-induced with 2% dextrose yeast samples were compared with the C) lupeol standard further D) co-injection analysis was performed to confirms the peak 1 is the product of the EutTTS2 gene

**Fig 5.**
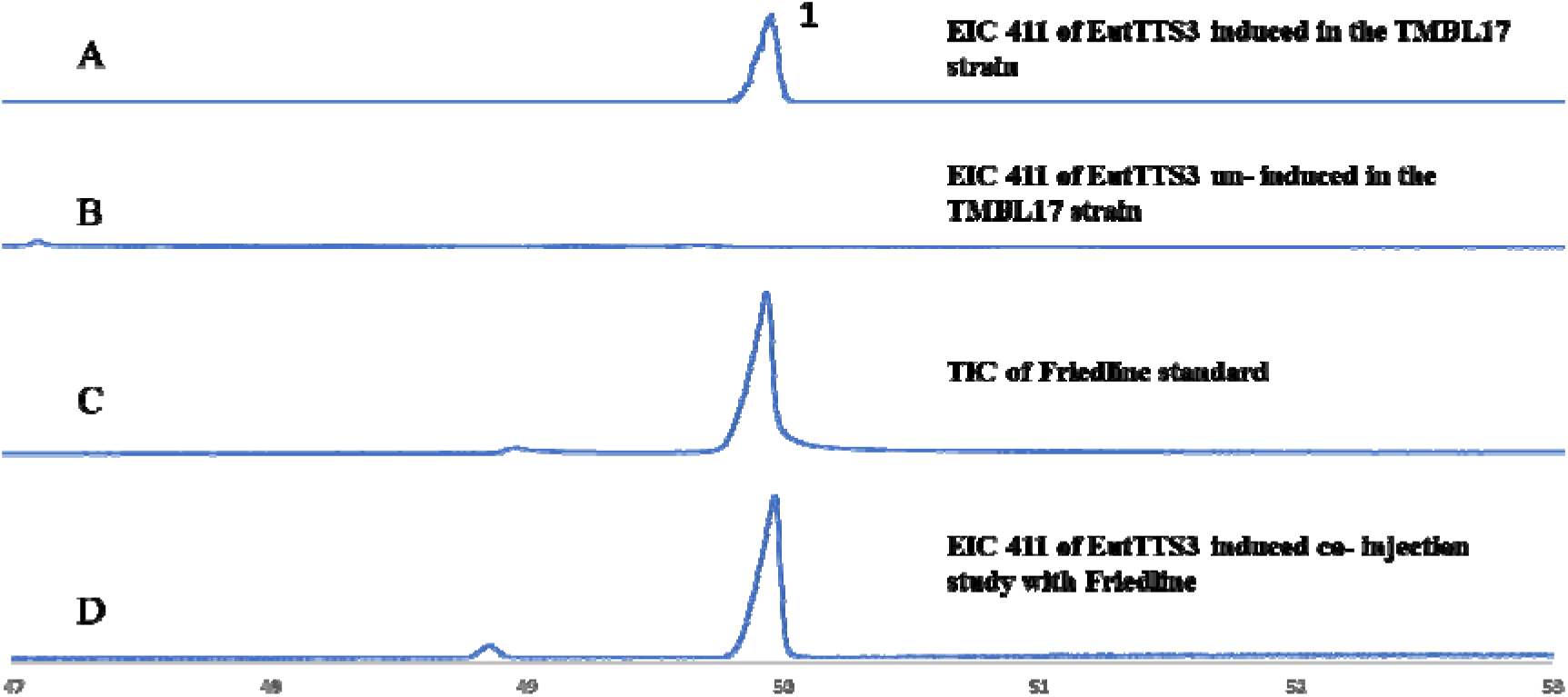
EIC (Extracted ion chromatogram) drawn from the TIC of total metabolites extracted from yeast (TMBL17 strain) harbouring EutTTS3 in pYES2/CT vector. EIC was obtained having the m/z value for 411 ion. A) On comparing EIC of total metabolites after induction with 2% galactose and B) un-induced with 2% dextrose yeast samples were compared with the C) friedline standard further D) co-injection analysis was performed to confirm the peak 1 is the product of the EutTTS3 gene.

**Fig 6.**
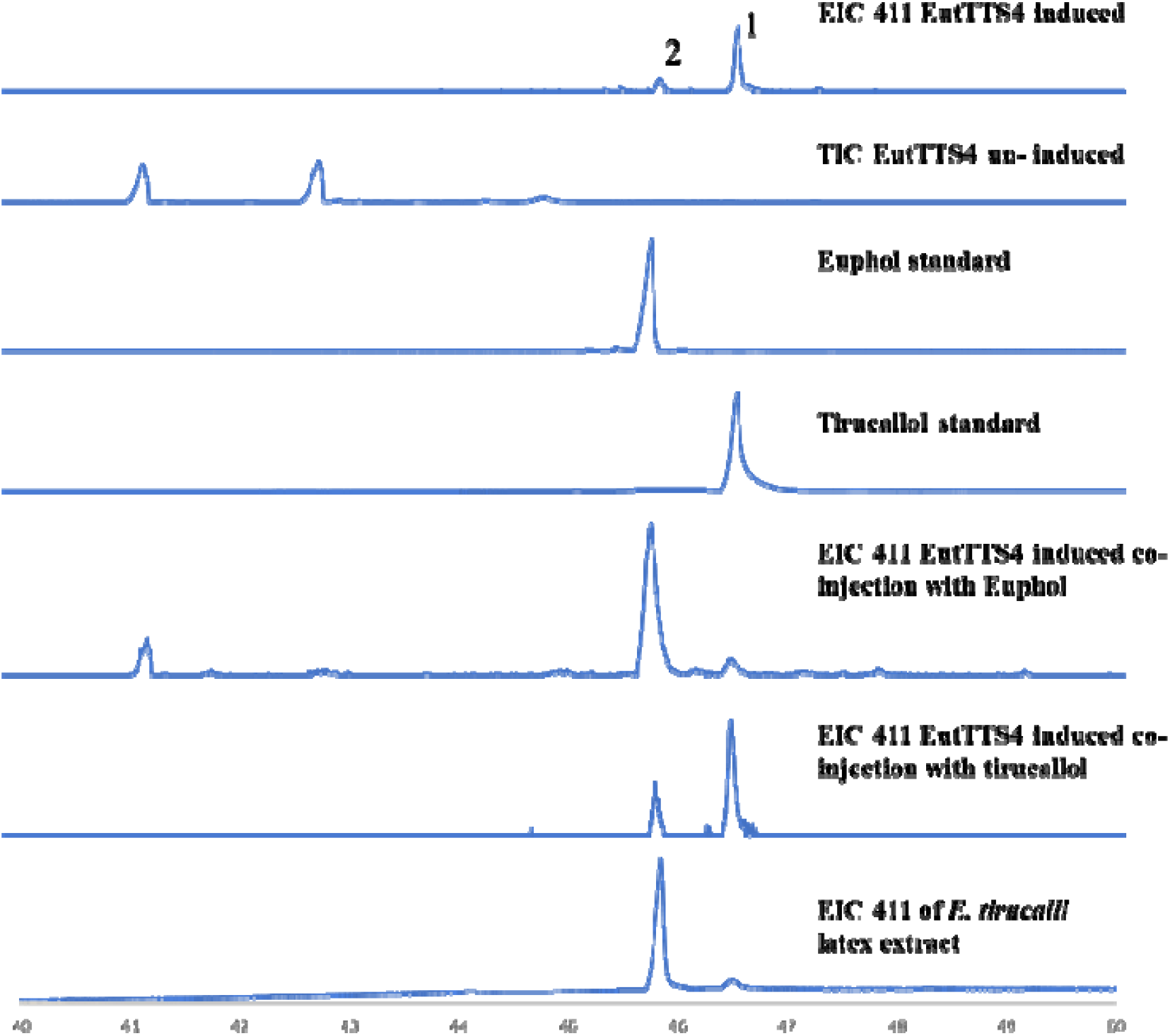
EIC (Extracted ion chromatogram) drawn from the TIC of total metabolites extracted from yeast (TMBL17 strain) harbouring EutTTS4 in pYES2/CT vector. EIC was obtained having the m/z value for 411 ion. A) On comparing EIC of total metabolites after induction with 2% galactose and B) un-induced with 2% dextrose yeast samples were compared with the C) Euphol standard and D) Tirucallol standard E) co-injection analysis with euphol and F) tirucallol was performed to confirm the peak 1 and 2 is the product of the EutTTS4 gene. G) Total metabolite extract from the latex of *E. tirucalli*.

**Fig 7.**
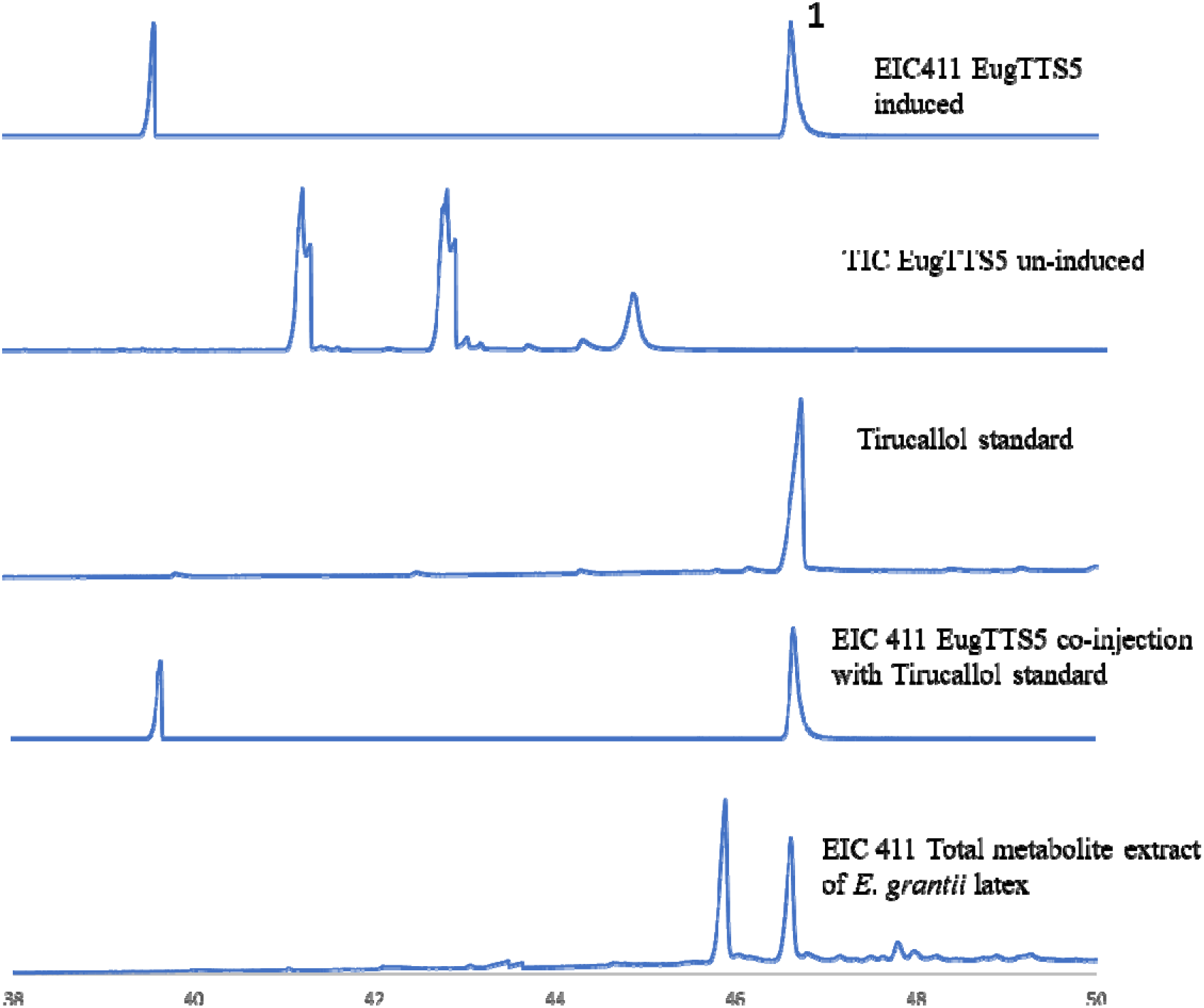
EIC (Extracted ion chromatogram) drawn from the TIC of total metabolites extracted from yeast (TMBL17 strain) harboring EugTTS5 in pYES2/CT vector. EIC was obtained having the m/z value for 411 ions. A) On comparing EIC of total metabolites after induction with 2% galactose and B) un-induced with 2% dextrose yeast samples were compared with the C) Tirucallol standard D) co-injection analysis with tirucallol was performed to confirm the peak 1 is the product of the EugTTS5 gene. E) TIC of total metabolites extracted from *E. grantii* latex.

### 2.6 Real-time analysis of HMGR and DXS

*E. grantii* and *E. tirucalli* are excellent sources of triterpenes, and latex is the primary source along with the different parts of the plant. Biosynthesis of triterpenes takes place from two different pathways Mevalonate and 2-C-methyl-D-erythritol-4-phosphate. As euphol and tirucallol are highly quantified triterpenes in *E. grantii* and *E. tirucalli,* to predict which pathway is mainly responsible for the biosynthesis of these two triterpenes. We selected HMGR (Hydroxymethyl glutaryl reductase) and DXS (D-xylulose synthase) from *E. tirucalli* and *E. grantii* (leaf and stem tissues). These two genes are the rate-limiting steps that mediate the metabolite flux of the MVA and MEP pathways. We designed the 200 base pair primers in real-time PCR against HMGR and DXS open reading frames (Table S5). The Ct value of HMGR was relatively low compared to DXS, as HMGR regulates the MVA pathway and DXS mediates the MEP pathway. The Low C_t_ value of HMGR indicates higher expression and confirms the involvement of the MVA pathway more in triterpene biosynthesis (Figure S5, S6).

### 2.7 Agroinfiltration of Euphol/Tirucallol synthases in *Nicotiana benthamiana*

To confirm the Euphol and tirucallol synthases transient transformation was carried out. The infiltration culture harboring EutTTS5 was optimized for infiltration studies using OD_600_ 0.2, 0.5 and 1. The abaxial side leaf of *N. benthamiana* was syringe infiltrated with the agrobacterium strain-GV3101 harboring EutTTS5 (Figure S20). The overexpression of EutTTS5 was observed in OD_600_ 0.2 infiltration culture quantified by real time PCR (Figure S21 A,B,C). whereas the metabolite analysis didn’t provide any significant result for EuTTS5. The EutTTS4 ORF was infiltrated into the *N. benthamiana* leaf, the overexpression was confirmed by real time PCR. Metabolite analysis of the transfected leaf’s has the euphol peak confirmed by the retention time, mass spectrum (Figure S19) and co-injection studies with the authentic standard (Figure S28).

### 2.8 Real time analysis of the triterpenes

Expression analysis of the triterpenes were performed by using the Roche Sybr green. All the five triterpenes functionally characterized were amplified with the primers spanning 200 bp and the amplification was estimated by using the Agilent Aria max real time PCR machine through SYBR green (Figure S25).

## 3.0 Materials and Method

### 3.1 Materials and chemicals

RNA was isolated using Spectrum™ Plant Total RNA Kit (Sigma- Aldrich) from the leaf ans stem tissues of *E. tirucalli* and *E. grantii* (Figure 1A). SuperScript® III First-Strand Synthesis System (Thermo Fisher Scientific, MA, USA) was used for cDNA synthesis. For PCR amplification, AccuPrime™ Pfx SuperMix (Invitrogen) and AccuTaq™ LA DNA Polymerase (Sigma-Aldrich) were used. For restriction digestion, New England BioLabs®inc (NEB) restriction enzymes were used. PCR product clean-up were carried out using PureLink™ PCR Purification Kit (Invitrogen). Ligation reaction was performed using T4 DNA ligases from Invitrogen and NEB. pESC-LEU (Agilent), pESC-TRP (Agilent) and pYES2/CT (Invitrogen) expression vectors were used. Cloning was carried out using Mach1™ T1R competent cells (Thermo Fisher Scientific). Yeast metabolite extracts were analyzed on Agilent 7890A GC coupled with 5975C mass detector.

### 3.2 Total RNA extraction from plant material

Leaf and stem tissues of *E. tirucalli* and *E. grantii* were collected from the NCL campus Pune. The leaf and stem tissues were collected in liquid nitrogen after washing with 0.1% DEPC water for molecular biology use. Total RNA was isolated from these tissues by Total RNA isolation kit, Sigma and the molecular integrity of RNA and the presence of DNA was checked by electrophoresis on the 1% DEPC agarose gel (Figure 1B). Further, Isolated RNA was treated with the DNase to remove the DNA contamination. Concentration and impurities of salt and proteins in the RNA samples were analyzed on Nanodrop (Thermo fisher). After analysis RNA from both the tissues ST- leaf and SB- the stem of *E. tirucalli* and LB- the stem, LL- leaf of *E. grantii* were sent for the next-generation sequencing.

### 3.2 Phytochemical analysis

*E. tirucalli* and *E. grantii* tissues leaf (5 g) and stem (5 g) tissue were crushed to powder under liquid nitrogen and extracted with ethyl acetate (10 mL × 3). The extraction of individual tissues was performed in triplicates with cholesterol as internal standard for the quantification purpose. The ethyl acetate layer was dried and concentrated under reduced pressure and analyzed on GC-MS. Ethyl acetate extract is being performed in GC-MS Agilent 7890A GC coupled with 5975C mass detector, Column Restek RTx-5ms (30m × 0.25mm × 0.25μm), 1.0 mL/min flow rate for the carrier gas helium. The program used for the analysis was 150 °C for 2 min followed gradient from 150°C to 250°C at 5°C/min for 11 minutes and finally temperature was maintained at 270°C for 15 min, injector inlet was maintained at 300 °C and detector at 280°C. 1 μl ethyl acetate extract was injected into the GC-MS RTx-5ms column. Compounds were identified and characterized by comparing the Rt value and mass spectrum with that of standards. Retention time of triterpenes present in the extract of ethyl acetate is (*R_t_* in min): 37.8 (tirucallol), 39.5 (euphol), 40.3 (β-amyrin), 42.0 (α-amyrin), 42.15 (lupeol), 42.4 (cycloartenol) and 47.1 (friedelin). The data were processed by MSD ChemStation Data Analysis (Agilent Technologies). The standard internal cholesterol was used in calculating the quantity of other terpenes by comparative TIC area analysis.

### 3.3 Transcriptome analysis

Total RNA was isolated from stem and leaf tissues of *E. tirucalli* and *E. grantii* (Figure 1A), mRNA sequencing libraries were prepared with Illumina-compatible Sure Select Strand-Specific RNA Library Prep Kit. Illumina NextSeq500 analysis generated an average 30 million reads for each tissue, which were trimmed by trimmomatic-0.32 (Bolger et al., 2014), a java-based package to remove the Illumina adapters along with the terminal sequences having Phred score lower than 33 (Table 1). Qualified reads were assembled using the Trinity-v2.6.6 package to generate the transcripts (Grabherr et al., 2011, Haas et al., 2013). Further, the cd-hit package generated 1,40,227 unigenes after removing the duplicate transcripts (Li and Godzik, 2006). After this 72.5 % of the unigenes were functionally annotated using the uniport database. Quality of unigenes assembly was checked using trinity pipeline to generate N10, N20, N30, N40, and N50 values. Unigenes of stem and leaf transcripts were subjected to virtual ribosomes- 2.0 generating the protein sequences in the six frames (Wernersson, 2006). Protein sequences with more than 20 amino acids were then submitted for the Pfam analysis in a 500 number pack on the Pfam- 35.0 server (Sonnhammer et al., 1997, Mistry et al., 2020). Transcripts with the Pfam_ID PF13243 PF13249 were identified and analyzed against the NCBI database using blastx to identify triterpene synthases. The ORF matching for triterpene synthase were shortlisted from *E. tirucalli* and *E. grantii* tissues and were subjected to the multiple sequence alignment (Clustal Omega) to remove the redundancy and generated thirteen unique triterpene synthases (Table 3). These raw reads were submitted on the SRA submission portal of NCBI (SAMN13519014/ST, SAMN13519015/SB, SAMN13519012/LL, and SAMN13519013/LB).

### 3.4 Phylogenetic analysis

Protein sequences of all functionally characterized triterpene synthase were downloaded from the UniProt database. Identified thirteen triterpene synthases from *E. tirucalli* and *E. grantii* were submitted to MEGA-X for the multiple sequence alignment by using muscle in FASTA format. Further, the alignment file was used to generate a phylogenetic tree using a neighbor-joining algorithm. The tree was saved in Newick format and modified in iTOL online software (Letunic and Bork, 2021).

### 3.5 CRISPR mediated knockout of erg7 and hem1 gene in Yeast (YPH499) strain

#### 3.5.1 Preparation of sgRNA for hem1 and erg7 genomic region and hem1 gene knock out in YPH499 (*S. cerevisiae*)

The open reading frame for genes which encodes hem1 and erg7 were retrieved from the NCBI database to identify the PAM sites with 5’ side 23 base pairs as gRNA in the form of two pair oligos (Table S7). pML107 has been used for the knockout of hem1 and erg7 genes in *S. cerevisiae,* this vector has gRNA cloning site BclI (methylated) and SwaI. To remove the methylation on restriction site BclI, this vector was transformed into the DNA cytosine methyltransferase and DNA adenine methyltransferase deficient *E. coli* cells purchased from the NEB. This transformation was modified at the step of secondary incubation, which was kept for 1 ½ hr. the unmethylated vector was isolated and gRNA for hem1 and erg7 was prepared as mentioned in (Laughery et al., 2015). Cloning of this gRNA of hem1 and erg7 were performed with the following reaction mixture, 1μL digested vector, and 1μL hybridized insert was kept ligation with 6μL Milli Q water, 1μL 10X NEB ligation buffer, and 1μL NEB ligase. This ligation reaction mixture was kept at 45⁰C for 5 minutes on ice immediately transformed into mach1 (Invitrogen) chemically competent cells by heat shock method. Cloning was confirmed by sequencing the plasmid with M13 forward and M13 reverse primers. After this Leucine2 cassette with promoter and terminator were amplified from the pESC-Leu vector by using primers with 40 bp overhangs identical to the genomic DNA of hem1 at 5’ and 3’ side from the chosen PAM site (Table S8). This amplicon was used as homologous directed repair (HDR) fragment. The gRNA containing vector pML107 was transformed into yeast YPH499 and the leucine2 HDR in a 1:4 ratio using the S.c easy comp Transformation kit. The transformation mixture was plated on leucine auxotrophic media containing 13 µg/mL hemin chloride and 20 µg/mL ergosterol. The positive colonies were confirmed using colony PCR with seq_hem1 forward and reverse primers (Table S7). Using following procedures;(Methods S2). The genome was isolated (Methods S1) and again checked for the presence of HDR fragment by PCR and then sent for the sanger sequencing. After confirming the knockout of hem1 in YPH499 yeast strain it was termed as TMBL17_1. Further, these colonies were again inoculated in 5ml YPD and made competent using S.c. Easy comp transformation kit and kept at −80°C.

#### 3.5.2 erg7 gene knock out using TMBL17_1 yeast strain

Similarly, as leu2 cassette, trp1 cassette was amplified from the pESC-Trp vector using primers having 40 bp overhangs identical to the genomic DNA of erg7 at the 5’ and 3’ side from the chosen PAM site (Table S7). pML107 containing gRNA fragment of erg7 along with trp1 cassette was transformed into TMBL1 using the S.c easy comp Transformation kit. Knock-out colonies were screened on CSM-w/o-leucine-tryptophan auxotrophic media having 13µg/mL hemin chloride and 20µg/mL ergosterol as an essential supplement. The colonies were analyzed using colony PCR method mention in (Methods S2) and using seq_erg7 forward and reverse primers (Table S8). The genome was isolated (Methods S1) again checked for the presence of HDR fragment by using seq_hem1 forward and reverse and seq_trp1 forward and reverse primers and then sent for the sequencing. After confirming the hem1 and erg7 knock out in yph499 we termed it TMBL17 yeast strain. Further, this knock-out yeast strain was grown in 500 ml YPD having 13µg/ml hemin chloride and 20µg/ml ergosterol as an external supplement for 3 days at 30 °C and 180 rpm. The pallet was harvested and processed as mentioned in (Pandreka et al., 2021) for metabolite analysis in GC-MS.

### 3.6 Heterologous expression and functional characterization of five triterpene synthases

Thirteen triterpene synthases were identified from the transcriptome analysis, and five were chosen for the functional characterization. The ORF of EutTTS1, EutTTS2, EutTTS3, EutTTS4, and EugTTS5 transcripts have an average length of 2.2 kb. Further, these transcripts were amplified and cloned in pYES2/CT yeast expression vector using the primers (Table S9). The cloned constructs of EutTTS1 and EutTTS2 were transformed in yeast INVSc-1 (*S. cerevisiae*), and construct EutTTS3, EutTTS4, and EugTTS5 were transformed in lanosterol deficient TMBL2 yeast cells (require external supplement 13 µg/mL hemin chloride and 20µg/ml ergosterol) by using *S. C.* Easy Competent cell preparation kit (Invitrogen). After this a single transformed colony was grown in CSM-w/o- ura media with 2 % glucose in 50 mL till O.D 600 reached 3.5. Induction was accomplished by transferring the culture into 2 % galactose containing CSM-w/o-ura 500 ml media at 30 □C and 180 rpm for 2 days. The pellet was harvested and processed by the method mentioned in (Pandreka et al., 2021). After that the extracts were dried and concentrated and analyzed in GC-MS.

### 3.7 Expression analysis of HMGR and DXS from *E. grantii* and *E. tirucalli*

Based on Pfam_ID, nine HMGR and three DXS have been identified from *E. grantii* and *E. tirucalli* leaf and stem tissues. These identified genes were subjected to expression analysis and 200-300 bp target region were chosen from these genes with the following primers (Table S5). The primers were designed as mentioned in (Pandreka et al., 2021). These primers have been used for the real-time PCR, which was having following components (1μL Forward primer, 1μL Reverse primer, 5μL Roche Sybr (2X), 1μL cDNA, and 2μLwater). This PCR was kept for the four different cDNA *E. grantii*, *E. tirucalli* leaf, and stem with three biological replicate and five technical replicates at 95 □C- 5 min_95 □C- 15 sec_54 □C- 1min. The generated C_t_ value was analyzed by the Livak method concerning the housekeeping gene Actin and EeF4.

### 3.8 Agroinfiltration of Euphol/tirucallol synthases in *Nicotiana benthamiana*

The involvement of EutTTS4 and EugTTS5 to produce euphol and tirucallol was also confirmed by the transient transformation in the leaf of *N. benthamiana*. ORF of both the transcripts was cloned in pRI101-AN at KpnI, SacI/SalI, KpnI restriction site by T_4_ DNA ligase from NEB (Table S9). pRI101 vector containing the gene was transformed into Agrobacterium tumefactions GV3101 chemical competent cells. A single transformed colony was inoculated in a 5 ml culture of LB medium plus appropriate antibiotics, and cells were grown overnight at 28°C and 180 rpm. This was sub-cultured in 50 ml LB and extended for another 16 hr at 28°C with the same rpm till OD_600_ reached 0.8. Agrobacterium was harvested by centrifugation at 4000 rpm and 4°C for 8 min. The cell pellet was diluted with MMA (10 mM MES, pH 5.6, 10 mM MgCl2, 100 mM acetosyringone) buffer till the cell density reached OD_600_ 0.2, 0.5, and 1. Further, it was incubated for 1 hr at 28 °C on an orbital shaker at 180 rpm. Immediately this culture was infiltrated into the greenhouse-grown, 4-5 weeks old *N. benthamiana* plants leaves infiltrated by pressing a 1 ml syringe against the leaf’s abaxial side and covering 75% of the leaf area. The experiment was performed with three replicates of empty vector, test, and control. After three days of incubation, the infiltrated leaf was harvested, and the total RNA was isolated using the Sigma (Total RNA isolation Spectrum kit). Total RNA was used for the cDNA synthesis using the First stand synthesis Superscript III kit, and the real-time PCR primers (Table S9) were optimized using JumpStart. The real-time PCR was kept with 1 μL cDNA and Roche sybr green by the program 95 °C- 30 sec, 95 °C- 15 sec, 54 °C- 1 min, 98- 30 sec on Agilent Ariamax. The harvested leaf was also crushed and resuspended in ethyl acetate for 1 hr at 180 rpm. The extract was dried and concentrated analyzed on GC-MS (Figure S41).

### 3.9 GC- MS analysis

1μL of total metabolite extracted from un- induced (control) and induced (test) samples of euphorbia triterpene synthase EuTTS1 and EuTTS2 were analyzed in GC-FID/GC-MS with HP-5MS capillary (30 m × 0.25 mm × 0.32 μm) columns. The injector was set at 250 ⁰C; the pressure was 9.3824 psi, septum purge flow was maintained at 3ml/min in splitless mode. The method used for the analysis was 80 ⁰C with a 2 min hold time. The temperature started rising at the rate of 5 ⁰C/min up to 290 ⁰C withhold time of 20 min. The whole program was of 64 minutes with 4 minutes of solvent delay. Product peaks were identified by injecting 1 mg/ml authentic triterpene standards (lupeol, Euphol, Tirucallol, β- amyrin, Friedelin, and α- amyrin).

## 4.0 Conclusion

In this study tissue, specific transcriptome analysis led us to identify thirteen unique triterpene synthases from *E. grantii* and *E. tirucalli*. Further, after blast analysis against the NCBI database, we found eight full-length triterpene synthases. Functional annotation of the eight-triterpene synthases was carried out. Based on blastx, phylogeny, and conserved amino acid analysis have identified two triterpenes EutTTS6 and EutTTS7 as β- amyrin synthase and EutTTS8 as cycloartenol synthase. The remaining five un-identified triterpene synthases were cloned and functionally characterized in yeast expression system; EutTTS1 as alpha amyrin synthase, EutTTS2 as lupeol synthase, EutTTS3 as friedelin synthase, and EutTTS4 as euphol and tirucallol synthase from *E. tirucalli.* EugTTS5 was characterized as tirucallol synthase from *E. grantii* whereas it is matching 98% with the euphol synthase it is having three key changes I666T, D667N and K701I which might be inhibiting the biosynthesis of euphol. Additionally, expression analyses of HMGR and DXS were performed to confirm the involvement of the MVA pathway more in triterpene biosynthesis than the MEP pathway. CRISPR-mediated knock-out of erg7 and hem1 genes and knock-in of leu3 and trp1 gene in yeast YPH499 strain was carried out to generate TMBL-17 yeast strain. In summary, this work adds two functionally new triterpene synthases and three new sequences of triterpene synthases in the triterpene biosynthetic pathway along with the highly efficient yeast strain to produce triterpene related products.

### Accession no

The accession number of these TTS are EutTTS1-MH880792.1, EutTTS2- MH880793.1, EutTTS3- MK387072.1, EutTTS4- MK387071.1, and EugTTS5- MK387070.1

## Acknowledgement

AK acknowledge CSIR, New Delhi for Fellowship. DM Acknowledge CSIR for Project funding. KT, AW, and SU acknowledge UGC, New Delhi for Fellowship. CSIR for Project funding MLP102426

## Data availability

All relevant data can be found within the published article and its supporting material.

## Supporting information

Additional supporting information may be found in the online version of this article

## Supporting information

Figure S1. Transcripts generated after assembly

Figure S2. Metabolite profiling in the four tissues of *E. tirucalli* and *E. grantii*

Figure S3. Fastqc analysis of the paired-end raw reads from the two tissues of *E. grantii* LB- stem, LL- leaf

Figure S4. Fastqc analysis of the paired-end raw reads from the two tissues of *E. tirucalli* SB- stem, ST- leaf.

Figure S5. DXS and HMGR qRT PCR identified from the *E. tirucalli* stem and leaf.

Figure S6. DXS and HMGR qRT PCR analysis identified from the *E. grantii* stem and leaf.

Figure S7. Showing the large-scale amplification of TTS. Lane 4, 5, 8, 18, and 21 is showing the 1 kb + ladder, lane 1, 7, 9, 17, and 20 is the control reaction. Lane 2, 6, 11, 16, and 19 is the amplification of EutTTS1, EutTTS2, EutTTS3, EutTTS4, and EugTTS5.

Figure S8. GC- MS chromatogram comparison with the α- amyrin standard and EutTTS1 induced and un- induced along with the co-injection analysis. The blue color represents the product peak of the test, and the red color represents un-induced, whereas 18HP0163 is of α -amyrin standard and 18HP0169 represents the co-injection study.

Figure S9. GC- MS chromatogram comparison with the lupeol standard and EutTTS2 induced and un-induced along with the co-injection analysis. The blue color peak represents the product peak of the test, and the red color represents the co-injection study with the lupeol standard, whereas the black color 20HP0560 is the peak of the lupeol standard and 20HP0567 is the un-induced chromatogram.

Figure S10. GC- MS chromatogram comparison with the Friedline standard and EutTTS3 induced and un-induced along with the co-injection analysis. The light green color peak represents the peak of the product, and the red one is the co-injection study, whereas the blue color represents the un-induced and black color peak represents the Friedline standard.

Figure S11. GC- MS chromatogram comparison with the tirucallol and euphol triterpene standards and EutTTS4 induced and un-induced along with the co-injection analysis. Red color peaks represent the product peaks of the gene black, and the light green peaks represent the co-injection study with euphol and tirucallol standards, whereas the blue color peaks represent the Euphol and tirucallol standard and the 220146.D represents the un-induced.

Figure S12. GC- MS chromatogram comparison with the tirucallol standard and EutTTS5 induced and un-induced along with the co-injection analysis. 210296.D represents the peak of the products biosynthesized from the gene and the red color represents the the co-injection study whereas the light green peaks represent the co-injection study and the blue color peaks shows the euphol and tirucallol standard.

Figure S13. GC- MS chromatogram comparison with the tirucallol and euphol triterpene standard and EutTTS4 transiently transformed into the *N. benthamiana* plantlets. The red color peak represents the product of transiently transformed EutTTS4, and the light green peak represents the co-injection study, whereas the blue color peak represents the euphol and tirucallol standard.

Figure S14. Mass spectrum of α- amyrin standard and the peak of EutTTS1.

Figure S15. Mass spectrum of lupeol standard and the peak of EutTTS2.

Figure S16. Mass spectrum of Friedline standard and the peak of EutTTS3.

Figure S17. Mass spectrum of tirucallol and euphol standard and the peak 1 and 2 of EutTTS4.

Figure S18. Mass spectrum of tirucallol standard and the peak of EutTTS5.

Figure S19. Mass spectrum of EutTTS5 peak and tirucallol standard.

Figure S20. Agrobacterium-mediated biotransformation of EutTTS4 (i2) and EutTTS5 (LB) cloned in pRI101 in *Nicotiana Benthamina* (tobacco plant).

Figure S21. Transient transformation expression analysis.

Figure S22. Amplification of the LEU2 HDR fragment and the colony PCR of 38 colonies with Seq_hem1 primers.

Figure S23. Amplification of the Trp1 cassette from the pESC-Trp and the colony PCR of 38 colonies with Seq_hem1 and Seq_erg7 primers.

Figure S24. Yeast genomic DNA isolated from hem1 mutant and hem1 and erg7 mutant colonies.

Figure S25. Real-time expression analysis of the triterpene synthase.

Figure S26. Quality check of assembled transcrits by N10-N50 values.

Figure S27. Active cite amino acid analysis of the triterpene synthases.

Figure S28. GC-MS chromatogram comparison of the EutTTS5 transient transformation in *N. benthamiana* along with the euphol triterpene standard and co-injection study. A) EIC411 of total metabolite analysis after transient. B) Transient transformation of pRI101-AN empty vector into the *Nicotiana benthamiana*. C) Euphol standard. D) Co-injection study of EutTTS5 product with Euphol standard.

Figure S29. Tritepene biosynthesis is derived fromo two different cations as intermediate.

Table S1. Triterpene synthases from the E. grantii stem tissue.

Table S2. Triterpene synthases identified from the E. grantii tissue leaf. Table S3. Triterpene synthases identified from the E. tirucalli stem.

Table S4. Triterpene synthases identified from the *E. tirucalli* leaf.

Table S5. primers for the real time PCR analysis of DXS and HMGR identified from E. grantii and E. tirucalli transcripts.

Table S6. Primers for the cloning of EutTTS and EugTTS transcript in pYES2/CT vector.

Table S7. Primers for the generation of gRNA and sequencing though the genome of TMBL-17.

Table S8. Primers for the sequencing of the vector constructs.

Table S9. Primers for the cloning of EutTTS4 and EugTTS5 transcript in pRI101 vector and for the real-time PCR analysis.

## Authors contribution

Ashish Kumar: Performed the transcriptome analysis, carried out the cloning and functional characterization of EutTTS1, EutTTS2, EutTTS3, EutTTS4 and EugTTS5, carried out the real time analysis and agroinfiltration study of EutTTS4 and EugTTS5, carried out the CRISPR-cas9 study to develop the TMBL17 lanosterol deficient yeast strain. Dhanashri S. Mulge: Involved in the development of TMBL17 yeast strain, involved in the functional characterization of EutTTS4 and EugTTS5, involved in real time PCR. Kalyani J. Thakar: Involved in the development of TMBL17 yeast strain, involved in real time PCR analysis. Avinash Pandreka: Involved in transcriptome analysis, involved in the functional characterization of EutTTS1 and EutTTS2. Amruta D. Warhekar: Involved in the Realtime PCR analysis. Sudha Ramkumar: Involved in the functional characterization of EutTTS2. Poojadevi Sharma: Carried out the quantification of Triterpene. Sindhuri Upadrasta: Involved in the CRISPR-cas9 experiment. Dhanasekaran Shanmugam: Involved in finalising and work flow of CRISPR-cas9, Hirekodathakallu V. Thulasiram: has conceptualized, supervised, and acted as overall study director

---

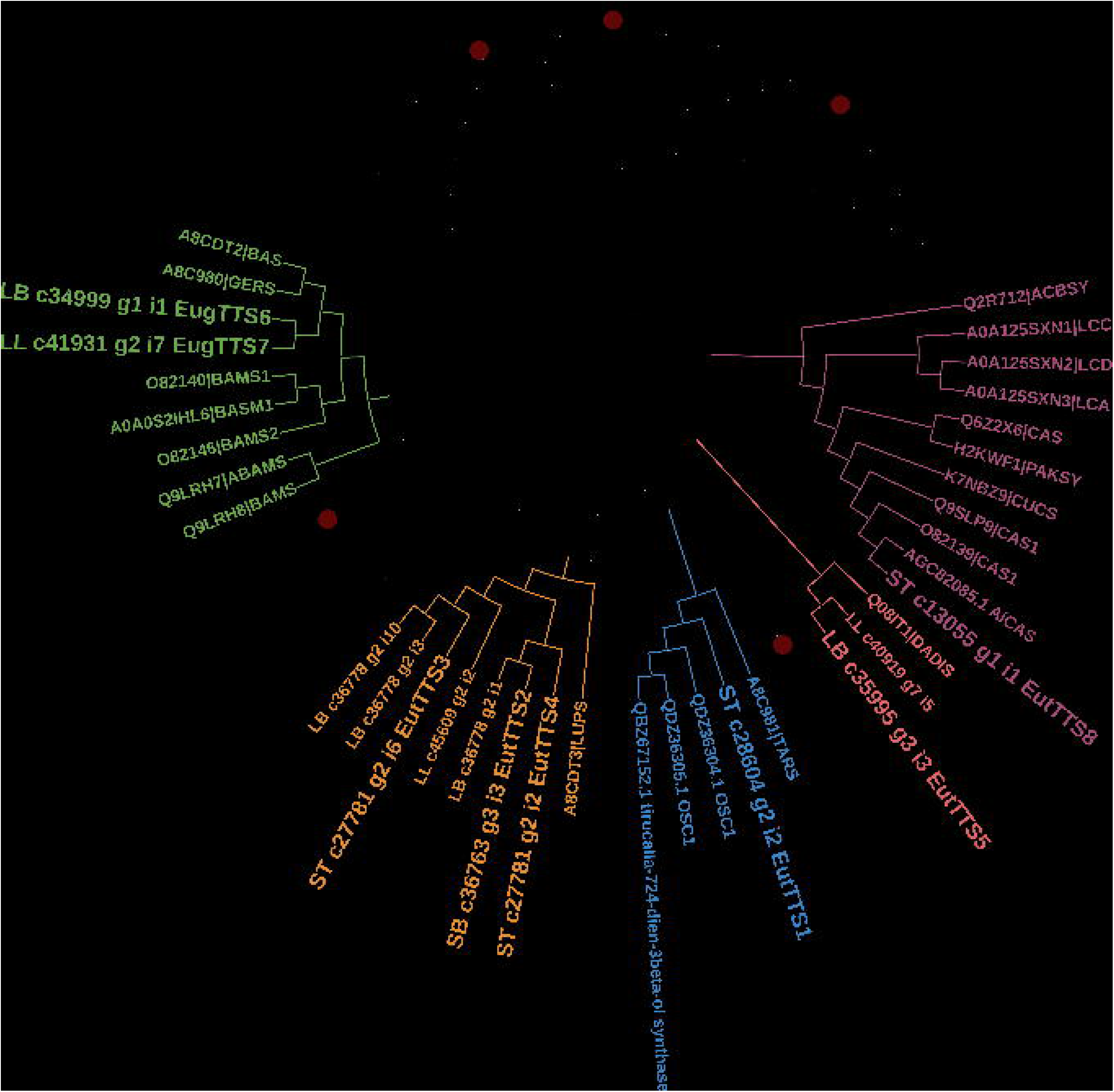

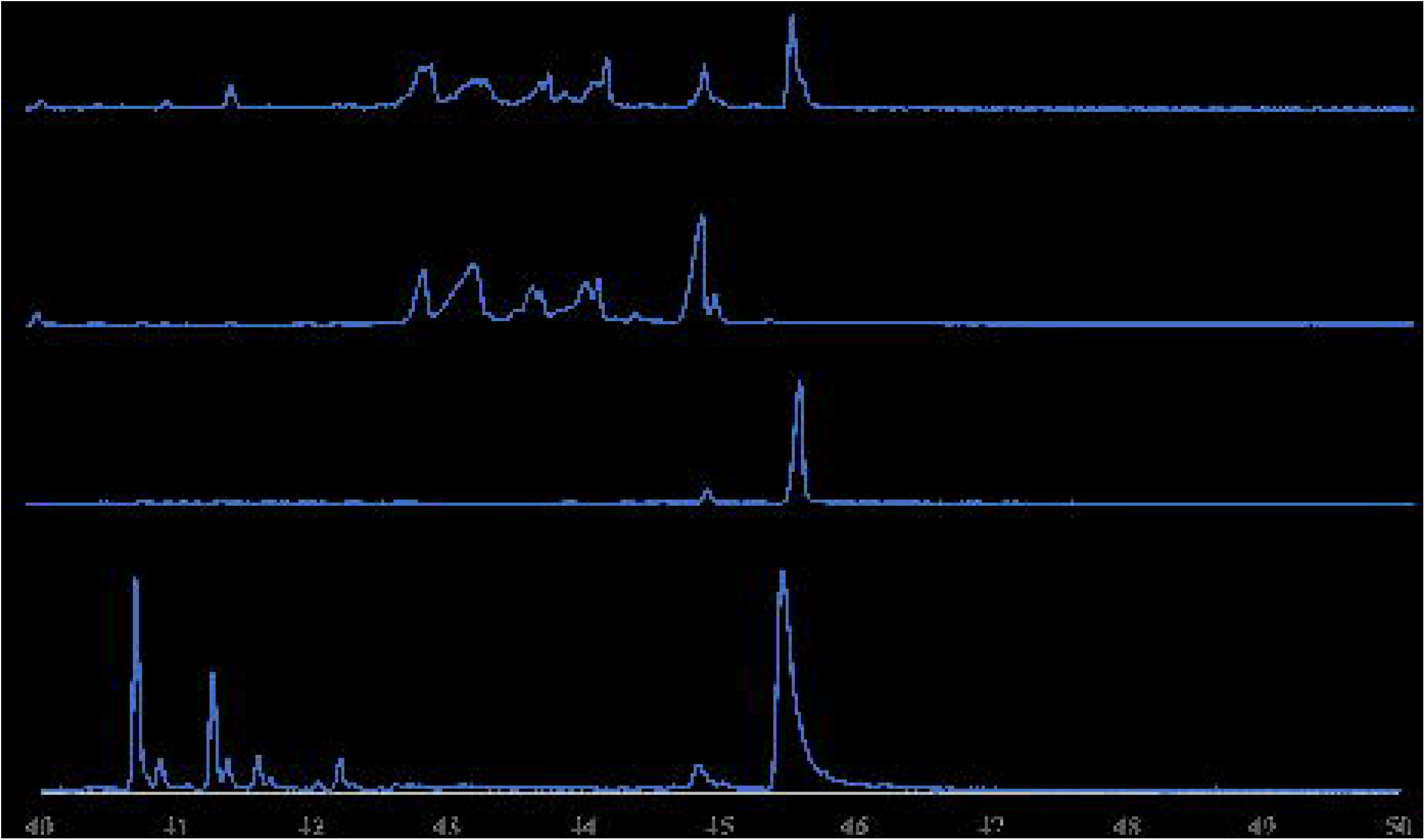

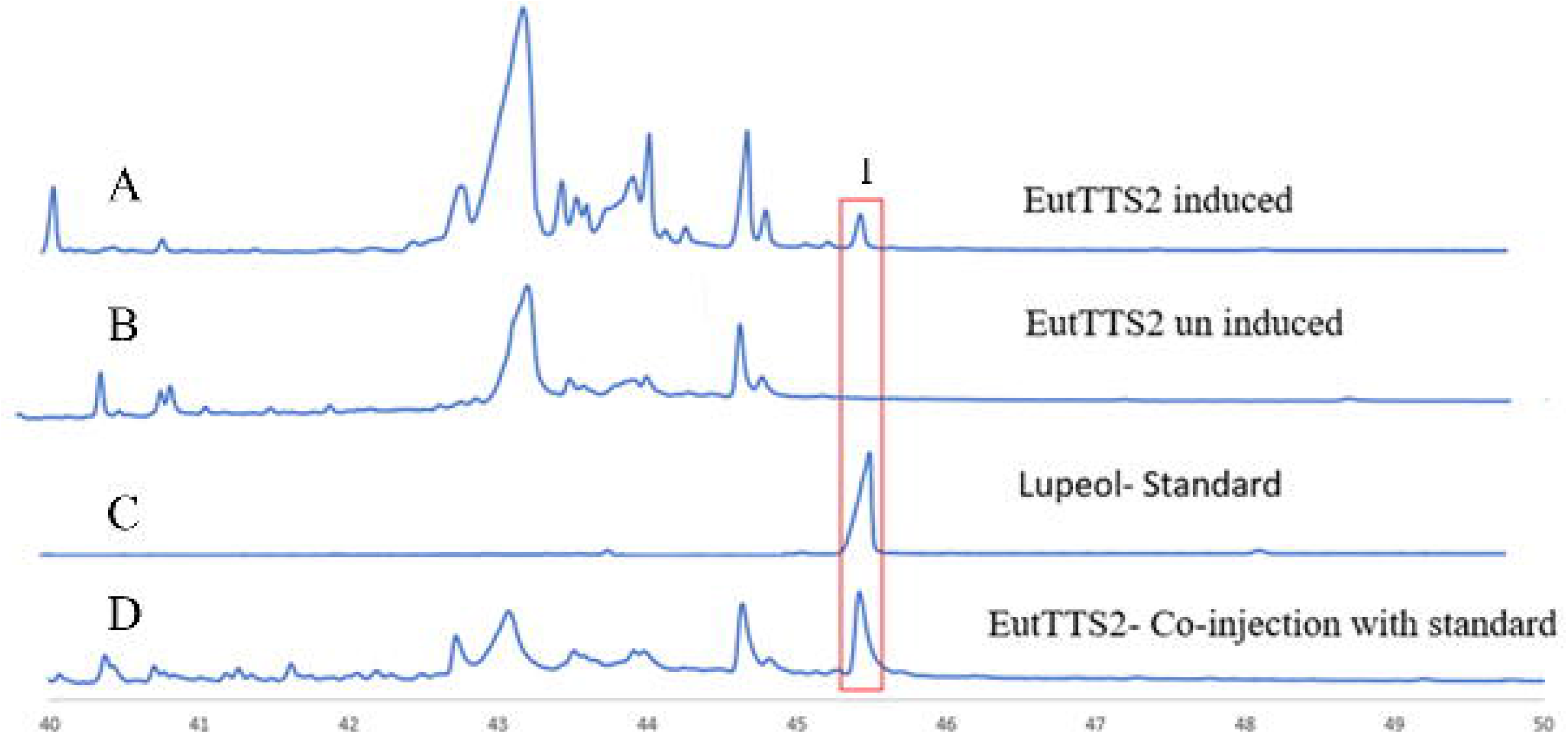

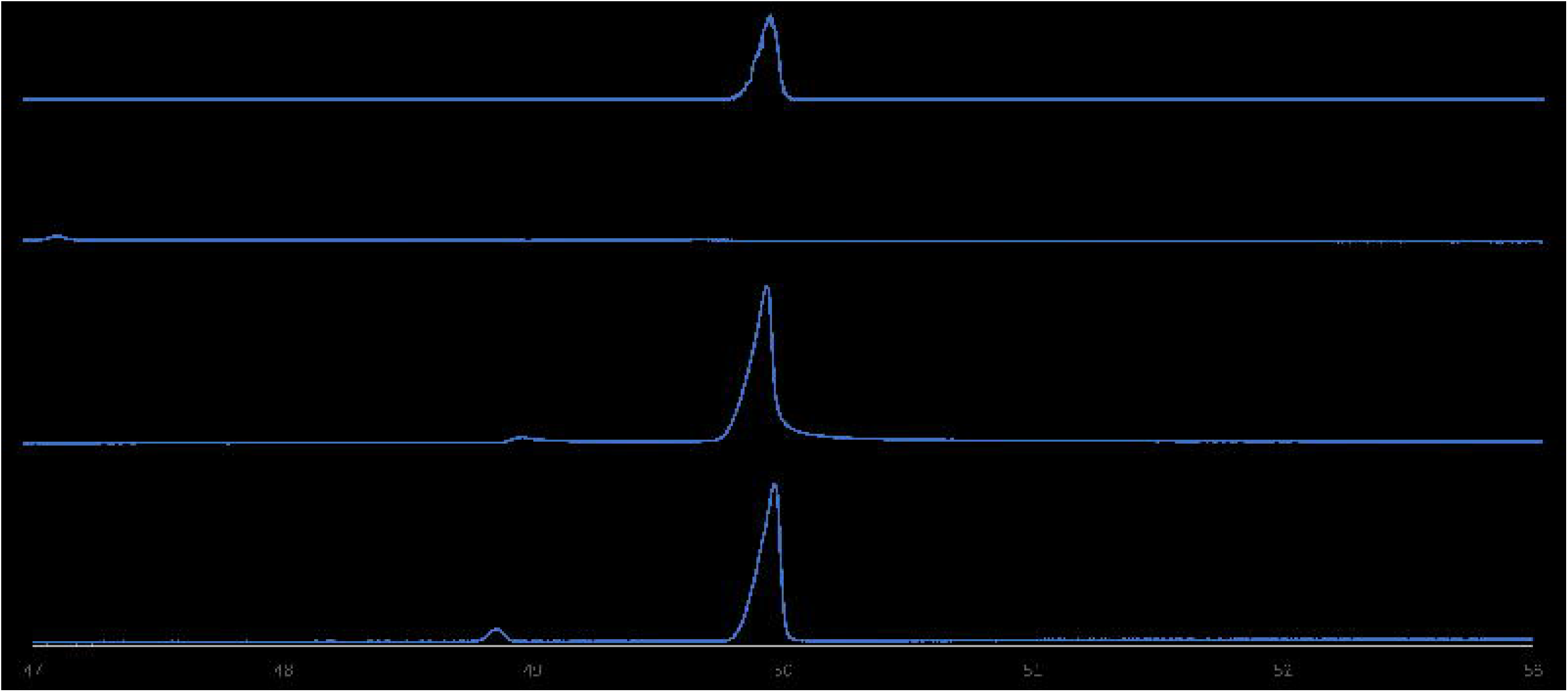

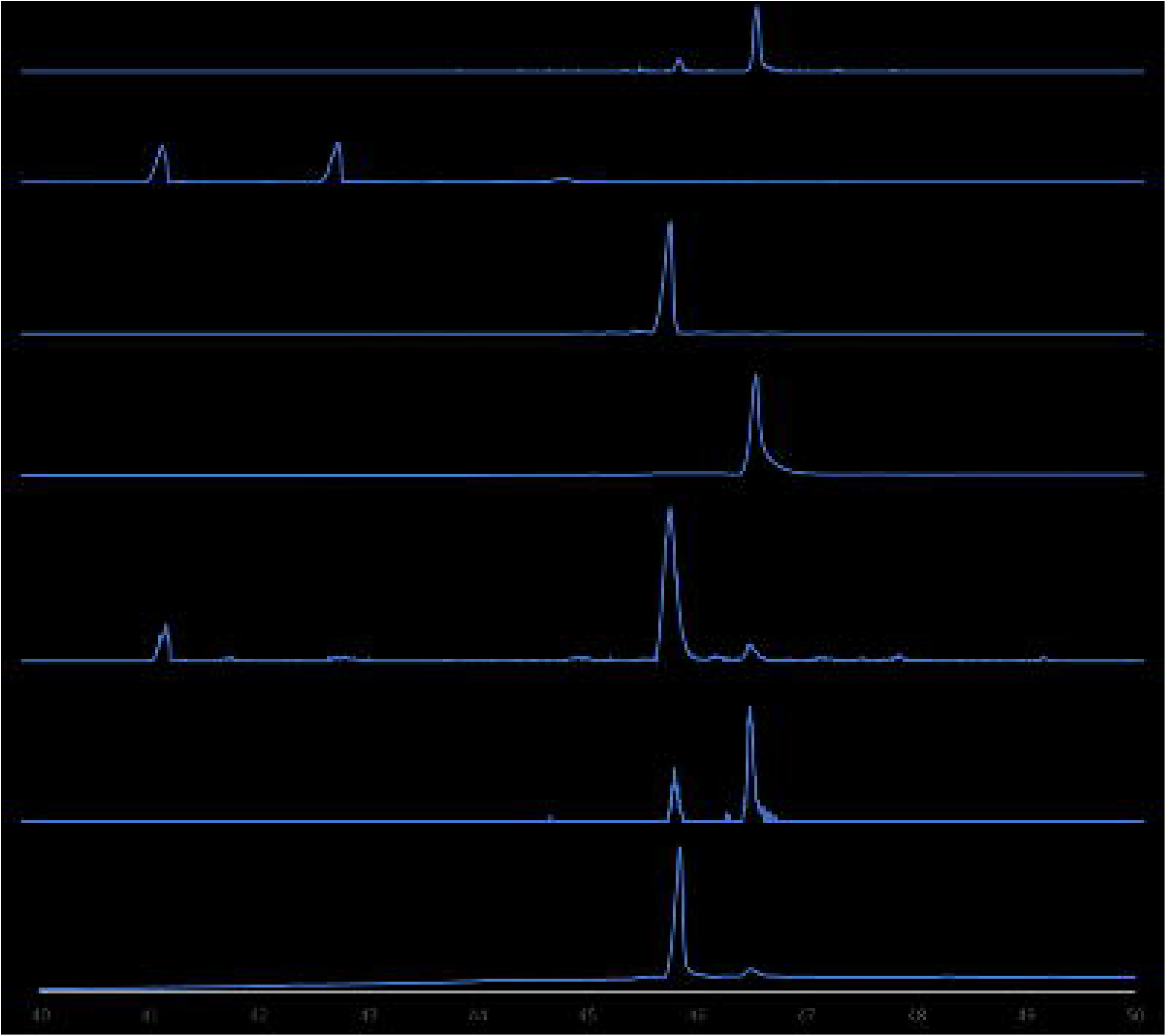

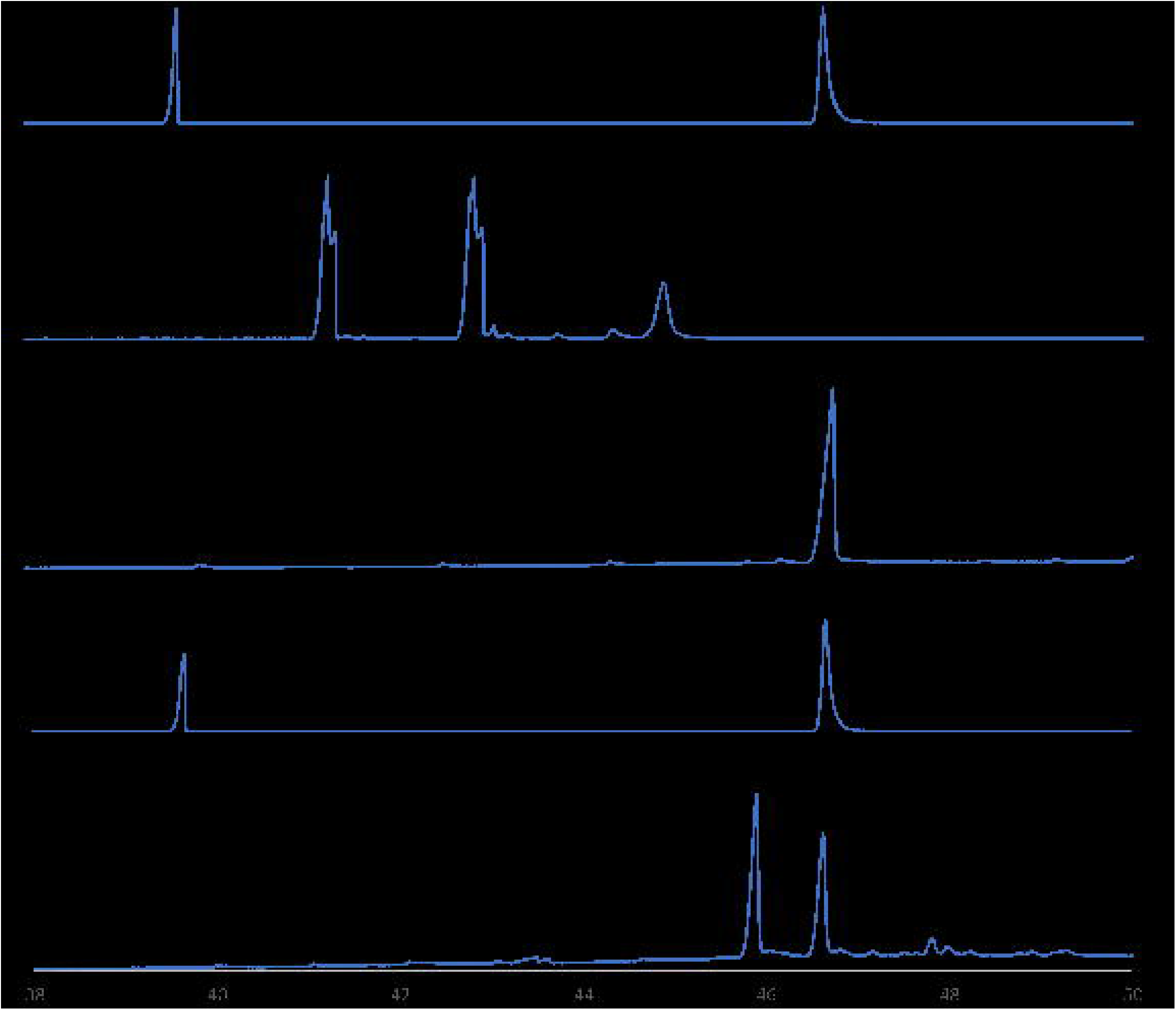

## Reference

Basyuni, M., Oku, H., Tsujimoto, E., Kinjo, K., Baba, S. & Takara, K. 2007. Triterpene synthases from the Okinawan mangrove tribe, Rhizophoraceae. FEBS J, 274, 5028-42. https://doi.org/10.1111/j.1742-4658.2007.06025.x

Bolger, A. M., Lohse, M. & Usadel, B. 2014. Trimmomatic: a flexible trimmer for Illumina sequence data. Bioinformatics (Oxford, England), 30, 2114-2120.

Corey, E. J., Cheng, H., Baker, C. H., Matsuda, S. P. T., Li, D. & Song, X. 1997. Studies on the Substrate Binding Segments and Catalytic Action of Lanosterol Synthase. Affinity Labeling with Carbocations Derived from Mechanism-Based Analogs of 2,3-Oxidosqualene and Site-Directed Mutagenesis Probes. Journal of the American Chemical Society, 119, 1289-1296.

Dale, M. P., Moses, T., Johnston, E. J. & Rosser, S. J. 2020. A systematic comparison of triterpenoid biosynthetic enzymes for the production of oleanolic acid in Saccharomyces cerevisiae. PLOS ONE, 15, e0231980.

Dang, T. & Prestwich, G. D. 2000. Site-directed mutagenesis of squalene–hopene cyclase: altered substrate specificity and product distribution. Chemistry & Biology, 7, 643-649.

Giner, J.-L. & Djerassi, C. 1995. A reinvestigation of the biosynthesis of lanosterol in Euphorbia lathyris. Phytochemistry, 39, 333-335.

Grabherr, M. G., Haas, B. J., Yassour, M., Levin, J. Z., Thompson, D. A., Amit, I., Adiconis, X., Fan, L., Raychowdhury, R., Zeng, Q., Chen, Z., Mauceli, E., Hacohen, N., Gnirke, A., Rhind, N., Di Palma, F., Birren, B. W., Nusbaum, C., Lindblad-Toh, K., Friedman, N. & Regev, A. 2011. Full-length transcriptome assembly from RNA-Seq data without a reference genome. Nature biotechnology, 29, 644-652.

Guhling, O., Hobl, B., Yeats, T. & Jetter, R. 2006. Cloning and characterization of a lupeol synthase involved in the synthesis of epicuticular wax crystals on stem and hypocotyl surfaces of Ricinus communis. Arch Biochem Biophys, 448, 60-72.

Haas, B. J., Papanicolaou, A., Yassour, M., Grabherr, M., Blood, P. D., Bowden, J., Couger, M. B., Eccles, D., Li, B., Lieber, M., Macmanes, M. D., Ott, M., Orvis, J., Pochet, N., Strozzi, F., Weeks, N., Westerman, R., William, T., Dewey, C. N, Henschel, R., Leduc, R. D, Friedman, N. & Regev, A. 2013. De novo transcript sequence reconstruction from RNA-seq using the Trinity platform for reference generation and analysis. Nature Protocols, 8, 1494-1512.

Hodgson, H., De La Peña, R., Stephenson, M. J., Thimmappa, R., Vincent, J. L, Sattely, E. S & Osbourn, A. 2019. Identification of key enzymes responsible for protolimonoid biosynthesis in plants: Opening the door to azadirachtin production. Proc Natl Acad Sci U S A, 116, 17096-17104.

Kajikawa, M., Yamato, K. T., Fukuzawa, H., Sakai, Y., Uchida, H. & Ohyama, K. 2005. Cloning and characterization of a cDNA encoding beta-amyrin synthase from petroleum plant Euphorbia tirucalli L. Phytochemistry, 66, 1759-66.

Kushiro, T., Shibuya, M., Masuda, K. & Ebizuka, Y. 2000. Mutational Studies on Triterpene Synthases:LJ Engineering Lupeol Synthase into β-Amyrin Synthase. Journal of the American Chemical Society, 122, 6816-6824.

Laughery, M. F., Hunter, T., Brown, A., Hoopes, J., Ostbye, T., Shumaker, T. & Wyrick, J. J. 2015. New vectors for simple and streamlined CRISPR-Cas9 genome editing in Saccharomyces cerevisiae. Yeast, 32, 711-20.

Letunic, I. & Bork, P. 2021. Interactive Tree Of Life (iTOL) v5: an online tool for phylogenetic tree display and annotation. Nucleic Acids Research, 49, W293-W296.

Li, W. & Godzik, A. 2006. Cd-hit: a fast program for clustering and comparing large sets of protein or nucleotide sequences. Bioinformatics, 22, 1658-9.

Liu, Y.-B., Yang, Y.-P., Yuan, H.-W., Li M.-J., Qiu, Y.-X., Choudhary, M. I. & Wang, W. 2018. A Review of Triterpenoids and Their Pharmacological Activities from Genus Kadsura. Digital Chinese Medicine, 1, 247-258.

Liu, Z., Zhang, Y., Sun, J., Huang, W.-C., Xue, C. & Mao, X. 2020. A Novel Soluble Squalene-Hopene Cyclase and Its Application in Efficient Synthesis of Hopene. Frontiers in Bioengineering and Biotechnology, 8.

Ma, L.-T., Wang, S.-Y., Tseng, Y.-H., Lee, Y.-R. & Chu, F.-H. 2013. Cloning and characterization of a 2,3-oxidosqualene cyclase from Eleutherococcus trifoliatus. Holzforschung, 67, 463-471.

Mistry, J., Chuguransky, S., Williams, L., Qureshi, M., Salazar, Gustavo A., Sonnhammer, E. L. L., Tosatto, S. C. E., Paladin, L., Raj, S., Richardson, L. J, Finn, R. D & Bateman, A. 2020. Pfam: The protein families database in 2021. Nucleic Acids Research, 49, D412-D419.

Niehaus, T. D., Kinison, S., Okada, S., Yeo, Y.-S., Bell, S. A, Cui, P., Devarenne, T. P & Chappell, J. 2012. Functional Identification of Triterpene Methyltransferases from Botryococcus brauniiRace B. Journal of Biological Chemistry, 287, 8163-8173.

Niehaus, T. D., Okada, S., Devarenne, T. P, Watt, D. S, Sviripa, V. & Chappell, J. 2011. Identification of unique mechanisms for triterpene biosynthesis in Botryococcus braunii. Proceedings of the National Academy of Sciences of the United States of America, 108, 12260-12265.

Ohyama, K., Suzuki, M., Kikuchi, J., Saito, K. & Muranaka, T. 2009. Dual biosynthetic pathways to phytosterol via cycloartenol and lanosterol in Arabidopsis. Proceedings of the National Academy of Sciences, 106, 725-730.

Pandreka, A., Chaya, P. S., Kumar, A., Aarthy, T., Mulani, F. A, Bhagyashree, D. D, B, S. H., Jennifer, C., Ponnusamy, S., Nagegowda, D. & Thulasiram, H. V. 2021. Limonoid biosynthesis 3: Functional characterization of crucial genes involved in neem limonoid biosynthesis. Phytochemistry, 184, 112669.

Qiao, W., Feng, W., Yang, L., Li, C., Qu, X. & Zhang, Y. 2021. De Novo Biosynthesis of the Anticancer Compound Euphol in Saccharomyces cerevisiae. ACS Synth Biol, 10, 2351-2358.

Sawai, S. & Saito, K. 2011. Triterpenoid Biosynthesis and Engineering in Plants. Frontiers in Plant Science, 2.

Sonnhammer, E. L., Eddy, S. R. & Durbin, R. 1997. Pfam: a comprehensive database of protein domain families based on seed alignments. Proteins, 28, 405-20.

Thulasiram, H. V., Erickson, H. K. & Poulter, C. D. 2007. Chimeras of Two Isoprenoid Synthases Catalyze All Four Coupling Reactions in Isoprenoid Biosynthesis. Science, 316, 73-76.

Thulasiram, H. V. & Poulter, C. D. 2006. Farnesyl Diphosphate Synthase:LJ The Art of Compromise between Substrate Selectivity and Stereoselectivity. Journal of the American Chemical Society, 128, 15819-15823.

Vernoud, V., Lebeigle, L., Munier, J., Marais, J., Sanchez, M., Pertuit, D., Rossin, N., Darchy, B., Aubert, G., Le Signor, C., Berdeaux, O., Lacaille-Dubois, M.-A. & Thompson, R. 2021. β-Amyrin Synthase1 Controls the Accumulation of the Major Saponins Present in Pea (Pisum sativum). Plant and Cell Physiology, 62, 784-797.

Wernersson, R. 2006. Virtual Ribosome--a comprehensive DNA translation tool with support for integration of sequence feature annotation. Nucleic Acids Res, 34, W385-8.

Zhou, J., Hu, T., Gao, L., Su, P., Zhang, Y., Zhao, Y., Chen, S., Tu, L., Song, Y., Wang, X., Huang, L. & Gao, W. 2019. Friedelane-type triterpene cyclase in celastrol biosynthesis from Tripterygium wilfordii and its application for triterpenes biosynthesis in yeast. New Phytologist, 223, 722-735.

