## Supplementary material for "Functional characterization of five triterpene synthases through De-novo assembly and transcriptome analysis of *Euphorbia grantii* and *Euphorbia tirucalli*": Figure S1: SI files revised 16-03-2023.docx

**Supplementary file**

A	B

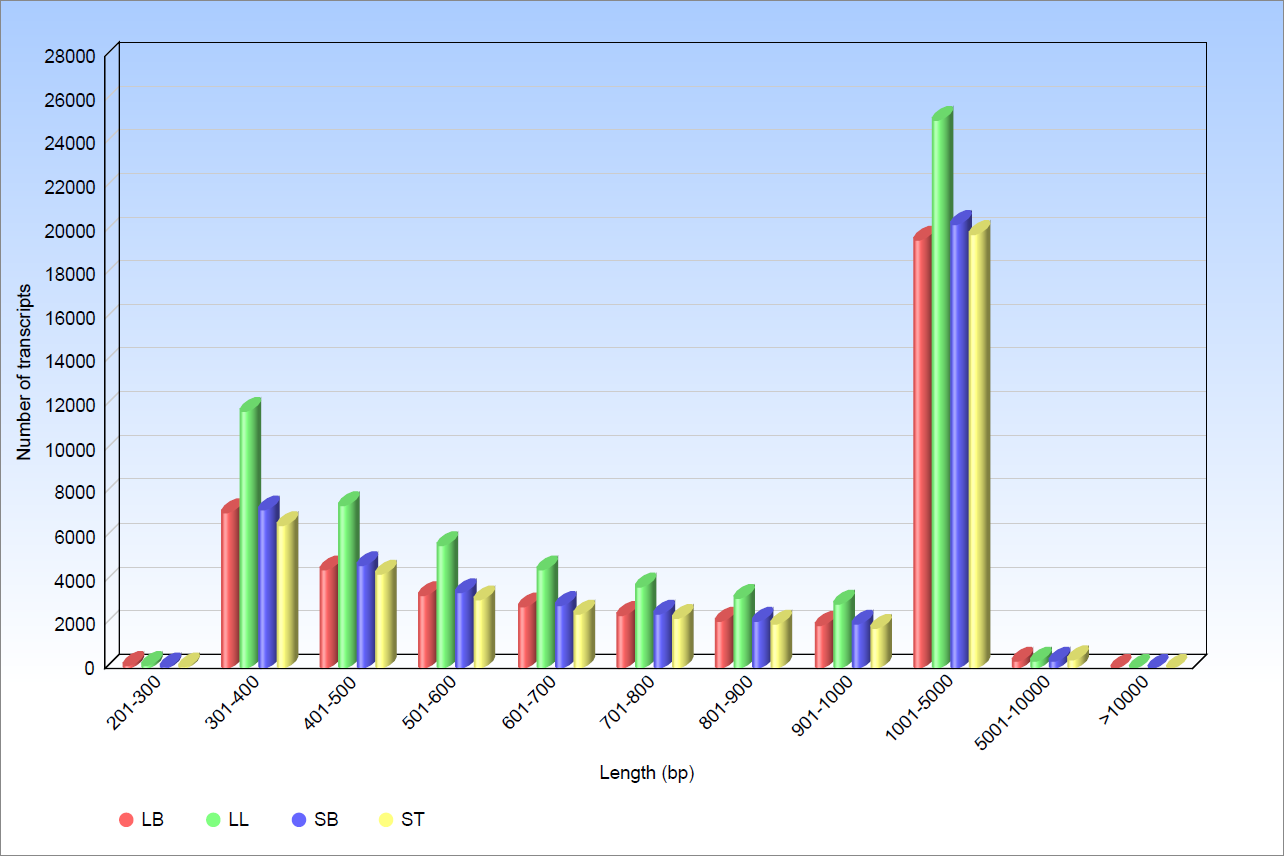

|  | KO_ID | Transcrpits |
| --- | --- | --- |
| LL | 6796 | 64694 |
| LB | 5781 | 44260 |
| SB | 5892 | 45665 |
| ST | 5971 | 42938 |

**Fig. S1** Transcripts generated after assembly (A) represent the length distribution of the various triterpenes. (B) KO_ID assigns to the total transcripts.

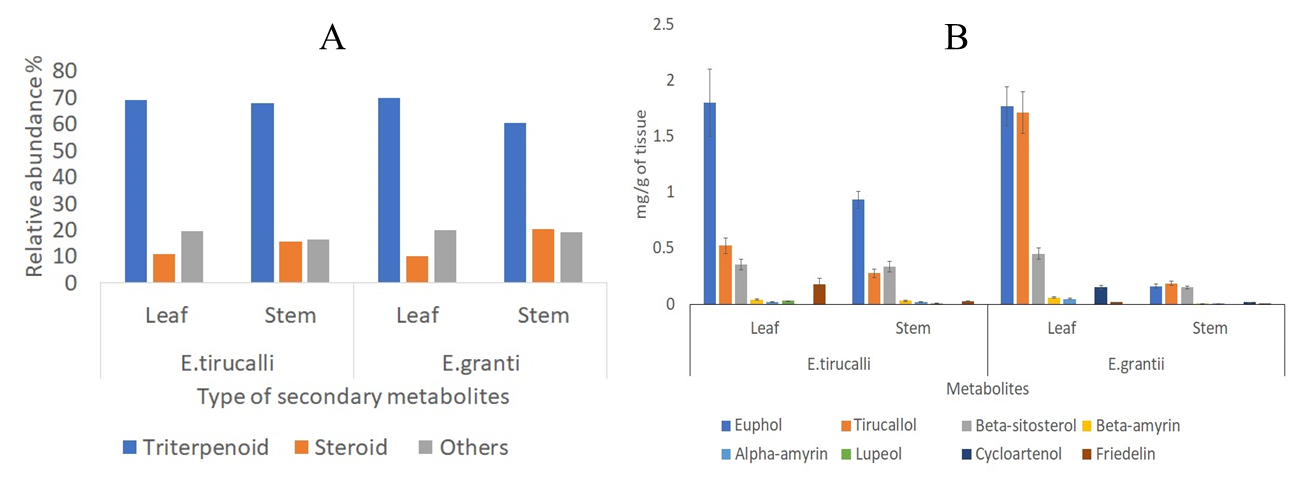

**Fig. S2** Metabolite profiling in the four tissues of *E. tirucalli* and *E. grantii.* (A) Total triterpenoid and steroid content in Euphorbia tirucalli and Euphorbia grantii leaf and stem tissue. (B)Amount of triterpenoid extracts obtained from stem and leaf tissue of *Euphorbia tirucalli* and *grantii*. Results represent the average of triplicates. *Cycloartenol was either absent or present in trace in *E. tirucalli* tissues. *Lupeol was either absent or present in trace in *E. grantii* tissues.

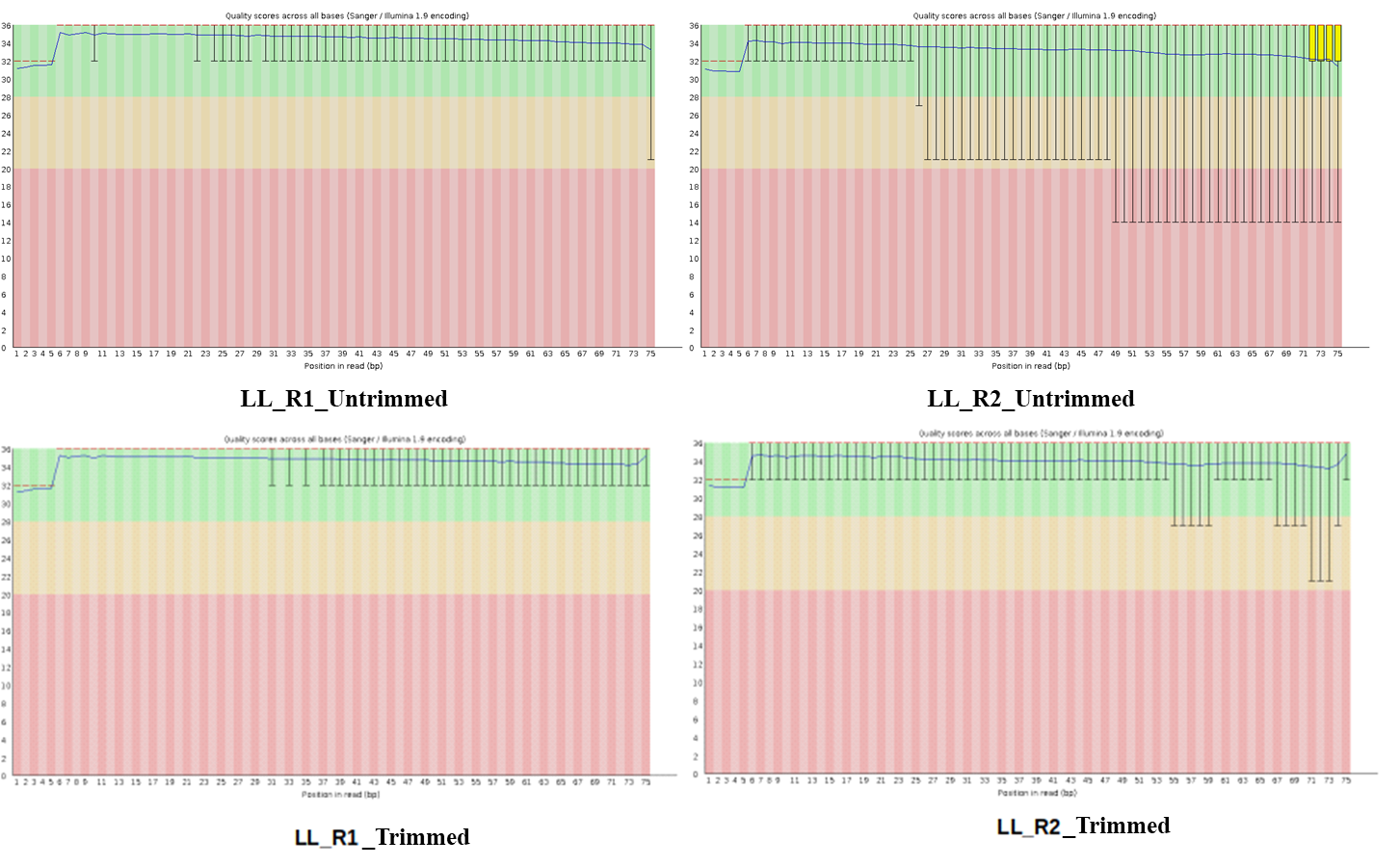

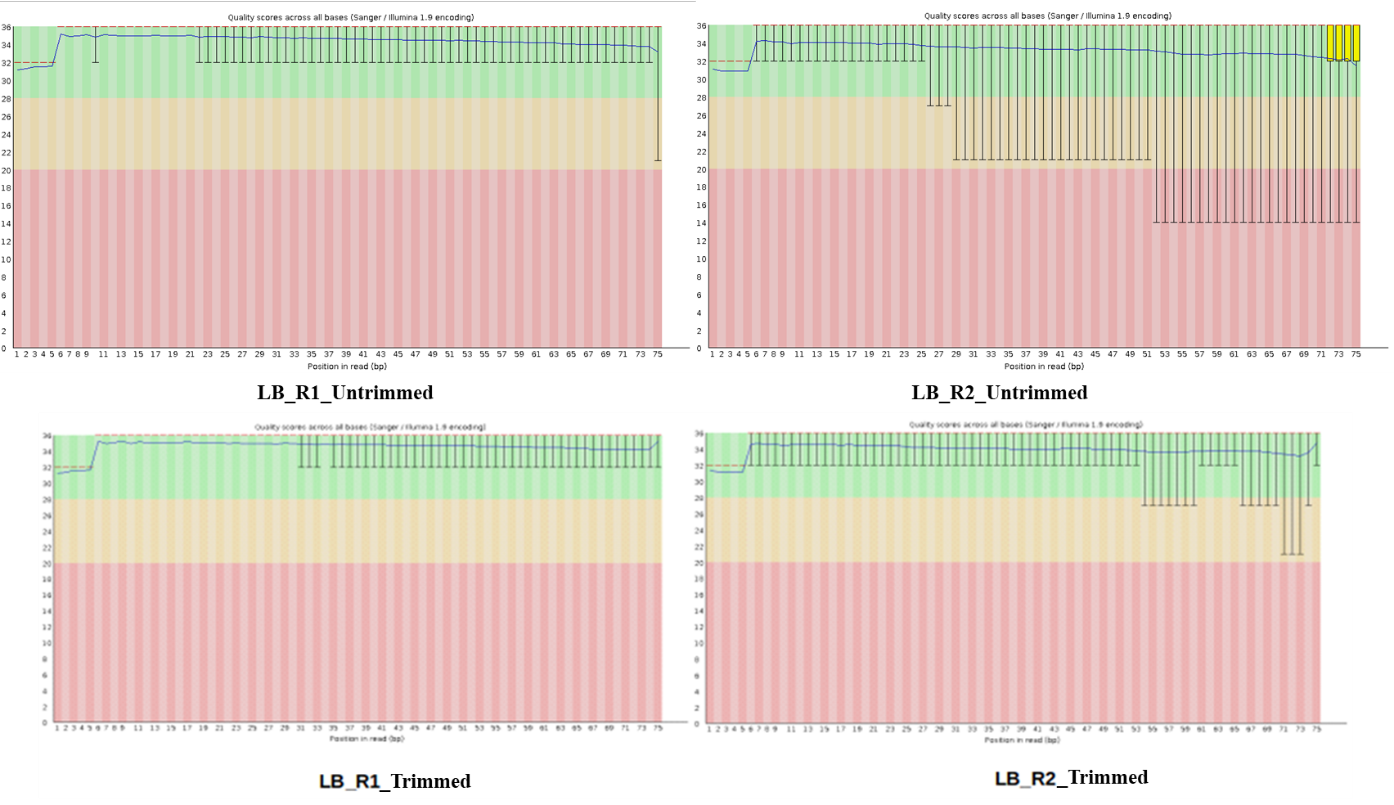

**Fig. S3** Fastqc analysis of the paired-end raw reads from the two tissues of *E. grantii* LB- stem, LL- leaf.

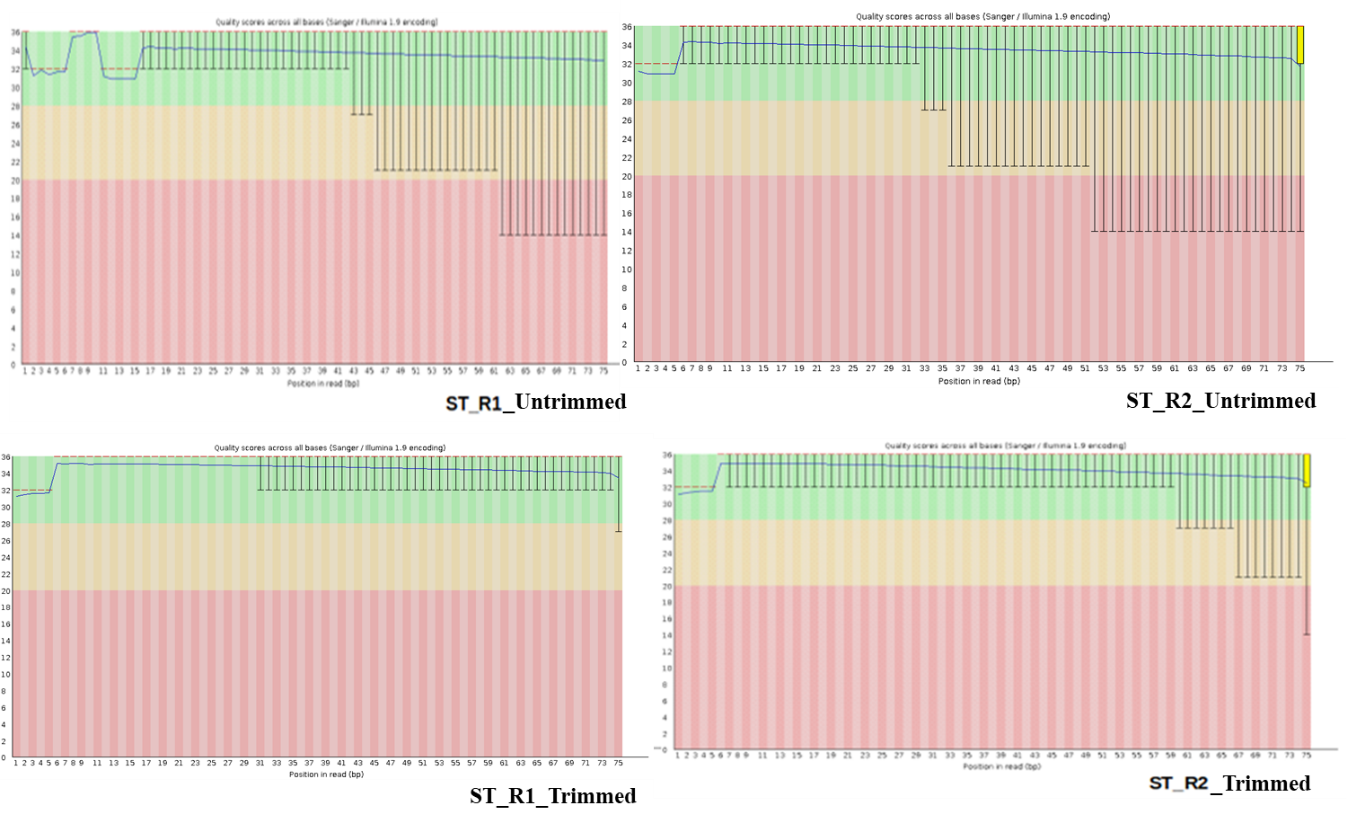

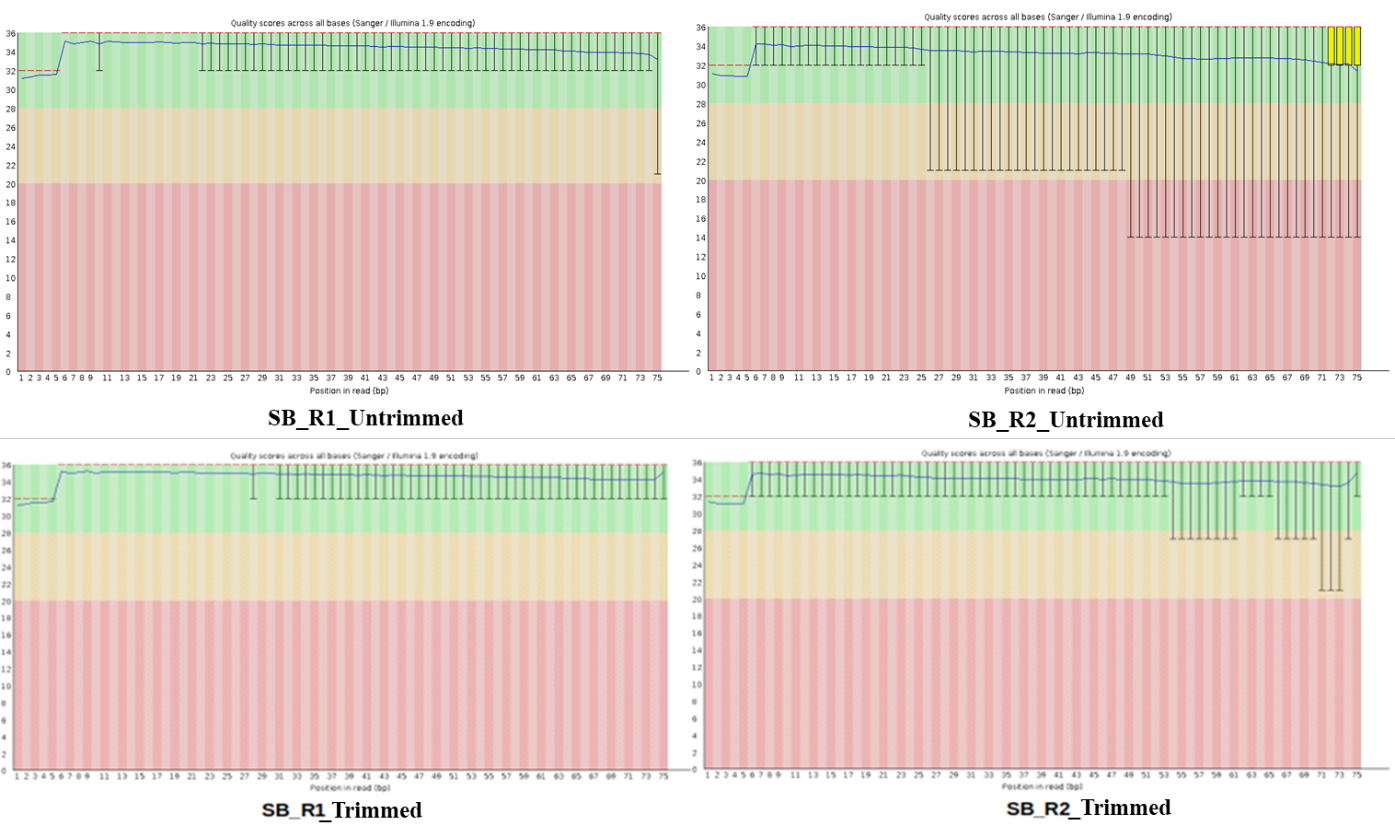

**Fig. S4** Fastqc analysis of the paired-end raw reads from the two tissues of *E. tirucalli* SB- stem, ST- leaf.

Euphorbia tirucalli

|  | leaf | Bark |
| --- | --- | --- |
| HMGCR1 | 1 | 0.0475673 |
| HMGCR2 | 0.0108369 | 0.0166234 |
| DXS | 4.56652e-05 | 0.00891648 |
| HMGCR3 | 2.71724e-05 | 0.00141692 |
| HMGCR4 | 3.47062 | 0.0156245 |

**Fig. S5** DXS and HMGR qRT PCR identified from the *E. tirucalli* stem and leaf.

Euphorbia grantii

|  | Leaf | stem |
| --- | --- | --- |
| HMGR1 | 1 | 194.74 |
| HMGR2 | 14.1656 | 6.00169 |
| HMGR3 | 0.293857 | 1.62859 |
| HMGR4 | 0.805248 | 0.0601172 |
| HMGR5 | 18.0864 | 21.5834 |
| DXS1 | 3.52681 | 0.00198997 |
| DXS | 1.21416 | 22.7325 |

**Fig. 6** DXS and HMGR qRT PCR analysis identified from the *E. grantii* stem and leaf.

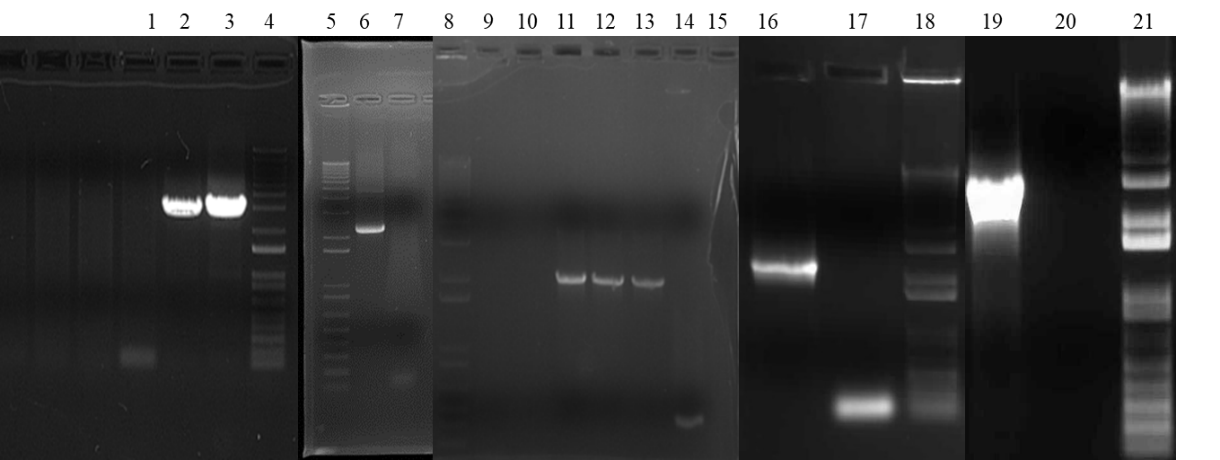

**Fig. S7** Showing the large-scale amplification of TTS. Lane 4, 5, 8, 18, and 21 is showing the 1 kb + ladder, lane 1, 7, 9, 17, and 20 is the control reaction. Lane 2, 6, 11, 16, and 19 is the amplification of EutTTS1, EutTTS2, EutTTS3, EutTTS4, and EugTTS5.

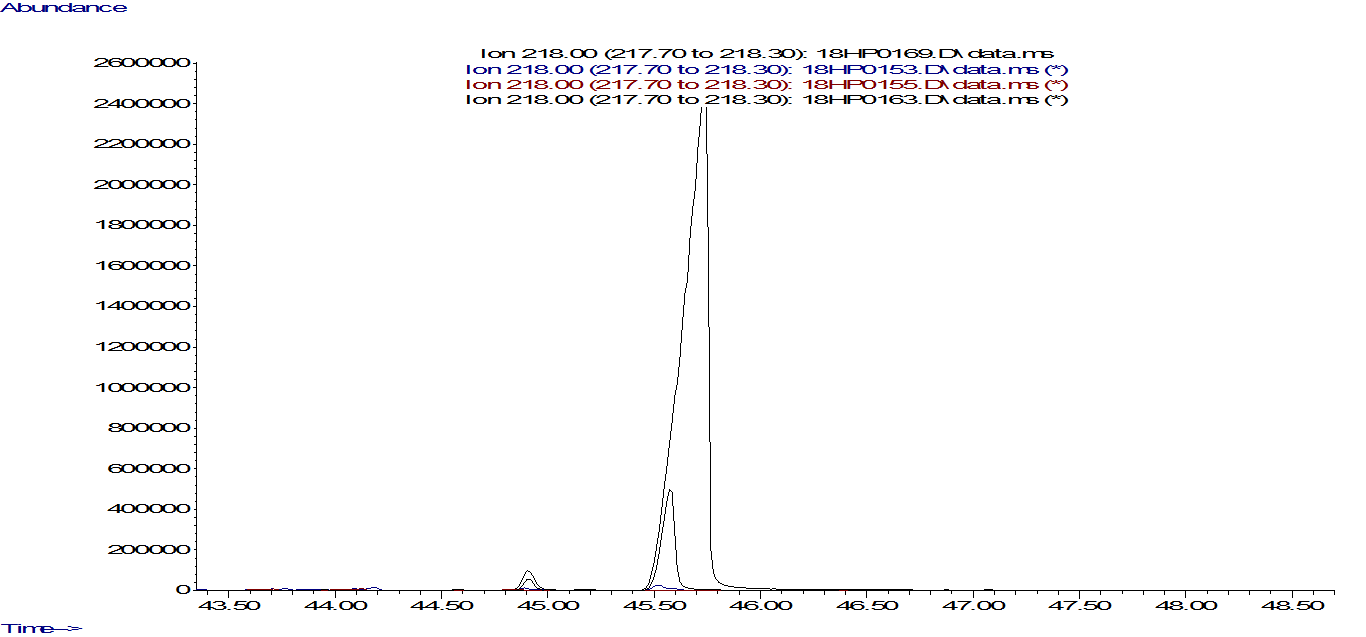

**Fig. S8** GC- MS chromatogram comparison with the α-amyrin standard and EutTTS1 induced and un- induced along with the co-injection analysis. The blue color represents the product peak of the test, and the red color represents un-induced, whereas 18HP0163 is of α-amyrin standard and 18HP0169 represents the co- injection study.

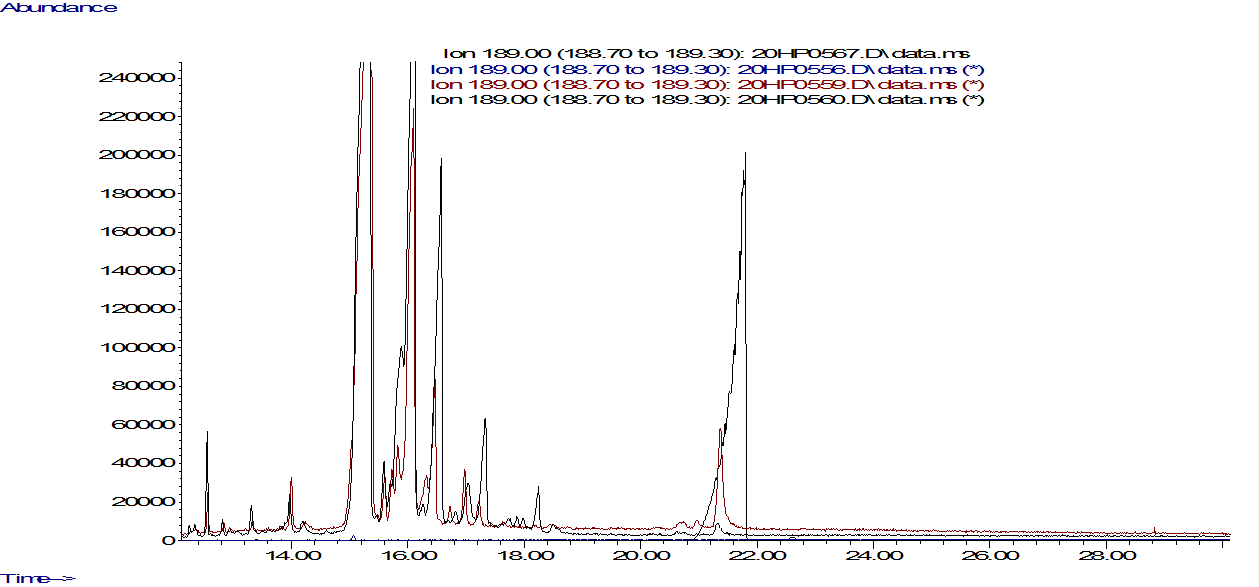

**Fig. S9** GC- MS chromatogram comparison with the lupeol standard and EutTTS2 induced and un-induced along with the co-injection analysis. The blue color peak represents the product peak of the test, and the red color represents the co-injection study with the lupeol standard, whereas the black color 20HP0560 is the peak of the lupeol standard and 20HP0567 is the un-induced chromatogram.

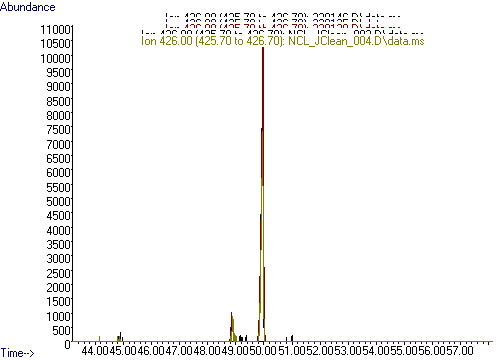

**Fig. S10** GC- MS chromatogram comparison with the Friedline standard and EutTTS3 induced and un-induced along with the co-injection analysis. The light green color peak represents the peak of the product, and the red one is the co-injection study, whereas the blue color represents the un-induced and black color peak represents the Friedline standard.

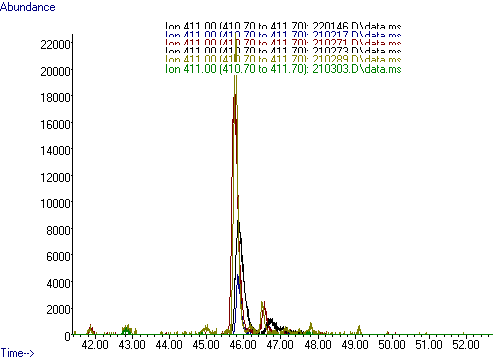

**Fig. S11** GC- MS chromatogram comparison with the tirucallol and euphol triterpene standards and EutTTS4 induced and un-induced along with the co-injection analysis. Red color peaks represent the product peaks of the gene black, and the light green peaks represent the co-injection study with euphol and tirucallol standards, whereas the blue color peaks represent the Euphol and tirucallol standard and the 220146.D represents the un-induced.

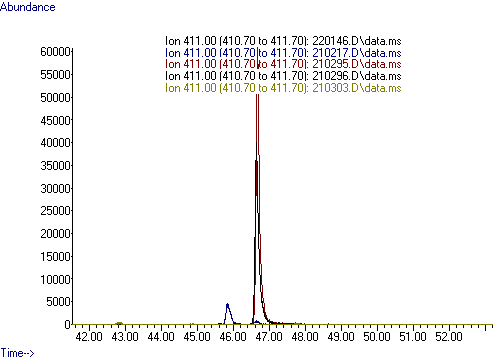

**Fig. S12** GC- MS chromatogram comparison with the tirucallol standard and EutTTS5 induced and un-induced along with the co-injection analysis. 210296.D represents the peak of the products biosynthesized from the gene and the red color represents the the co- injection study whereas the light green peaks represents the the co- injection study and the blue color peaks shows the euphol and tirucallol standard.

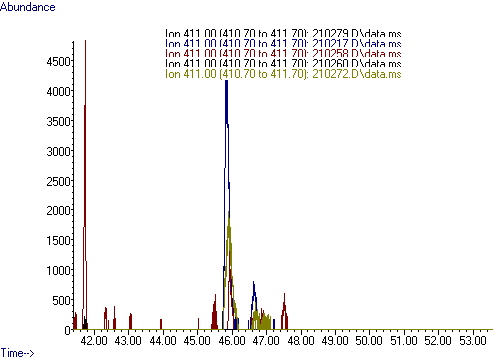

Fig. S13 GC- MS chromatogram comparison with the tirucallol and euphol triterpene standards and EutTTS4 transiently transformed into the *N. benthamiana* plantlets. The red color peak represents the product of transiently transformed EutTTS4, and the light green peak represents the co-injection study, whereas the blue color peak represents the euphol and tirucallol standard. 210260.D is the empty vector chromatogram.

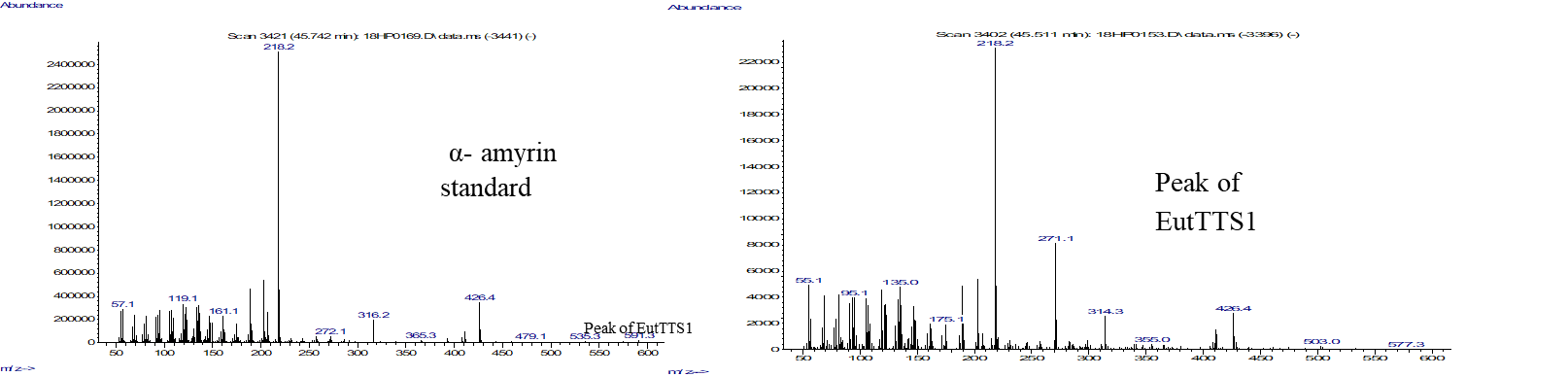

**Fig. S14** Mass spectrum of α-amyrin standard and the peak of EutTTS1.

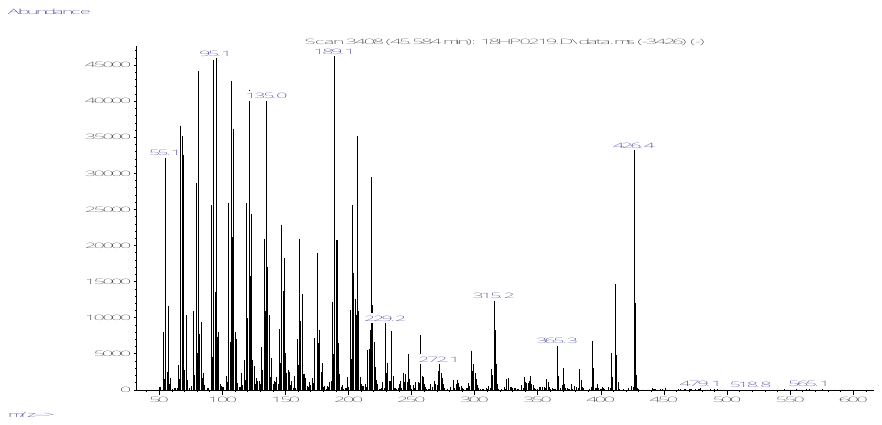

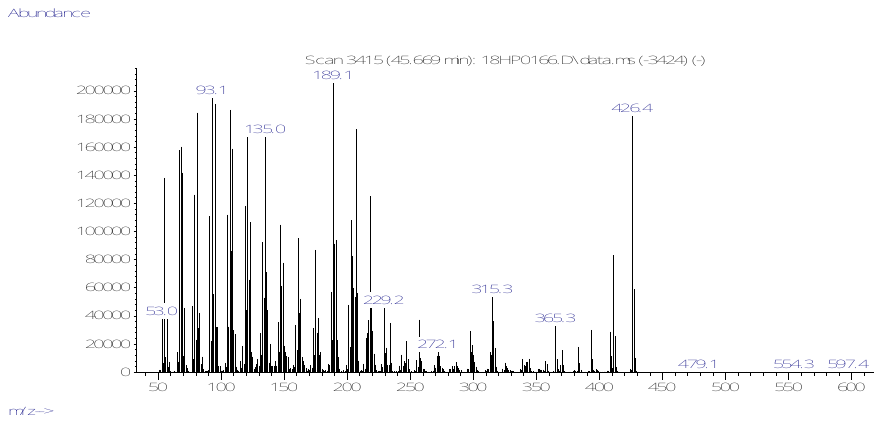

Peak of EutTTS2

Lupeol Standard

**Fig. S15** Mass spectrum of lupeol standard and the peak of EutTTS2.

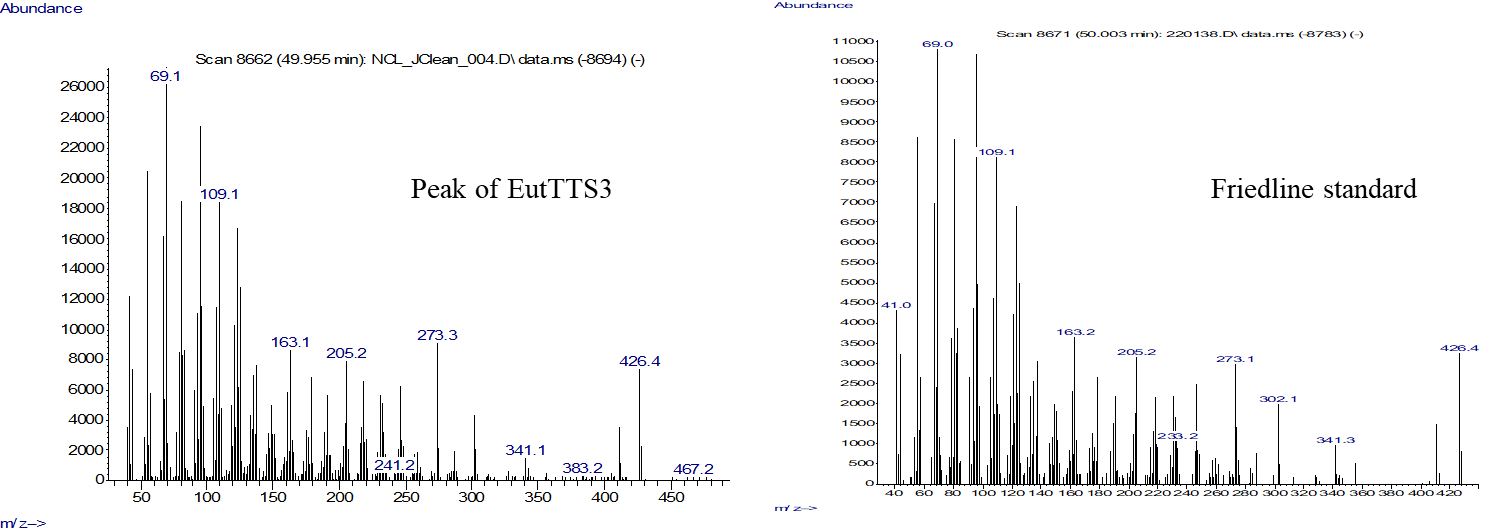

**Fig. S16** Mass spectrum of Friedline standard and the peak of EutTTS3.

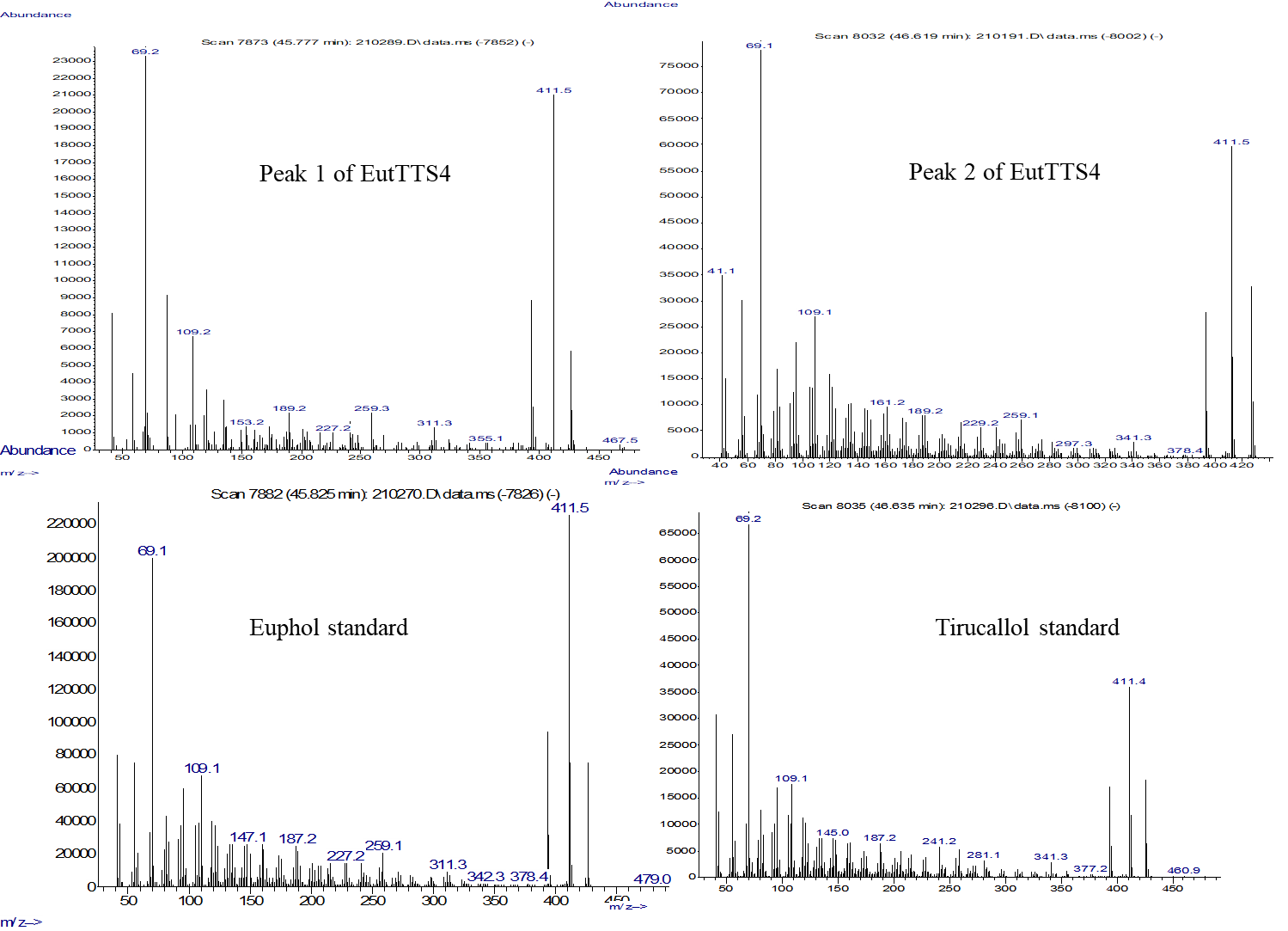

**Fig. S17** Mass spectrum of tirucallol and euphol standard and the peak 1 and 2 of EutTTS4.

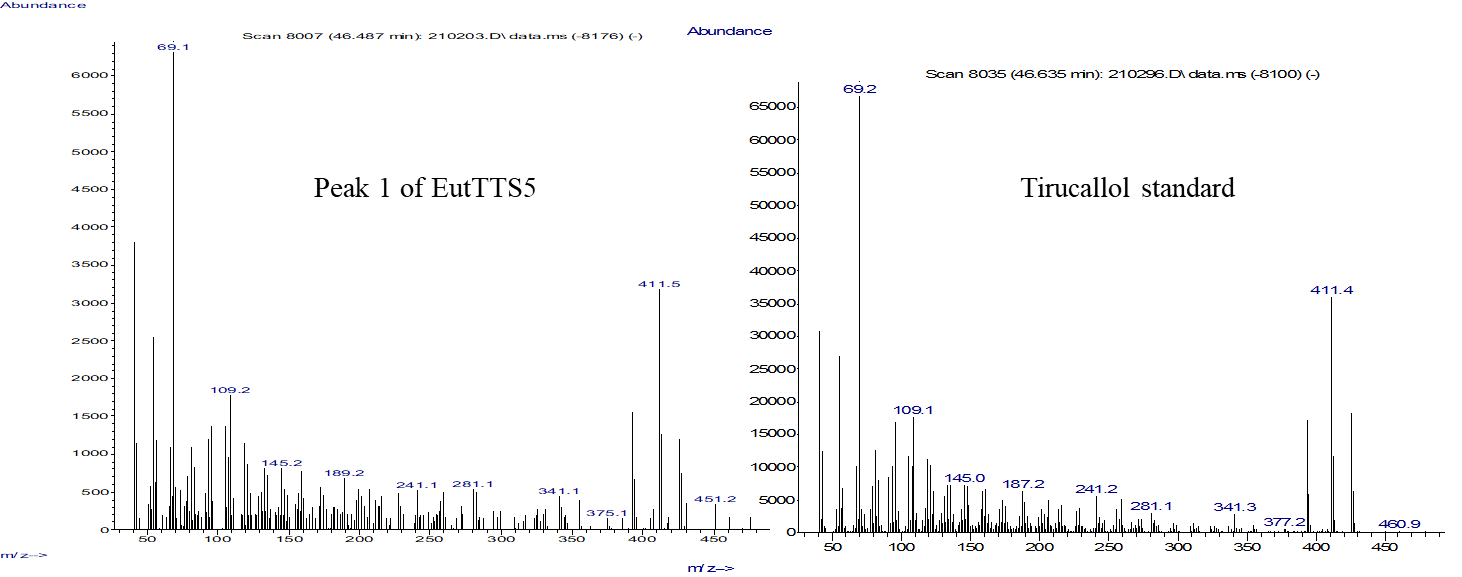

**Fig. S18** Mass spectrum of tirucallol standard and the peak of EutTTS5.

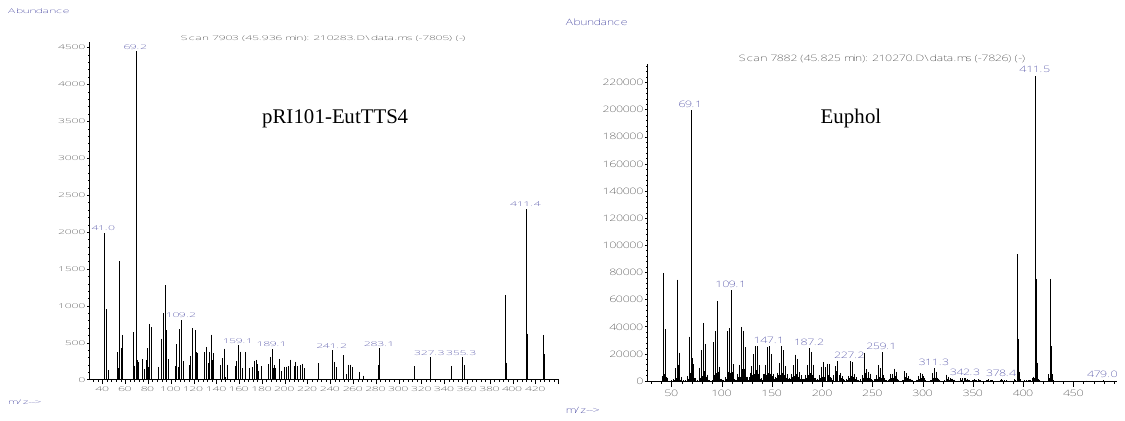

**Fig. S19** Mass spectrum of EutTTS4 peak and euphol standard.

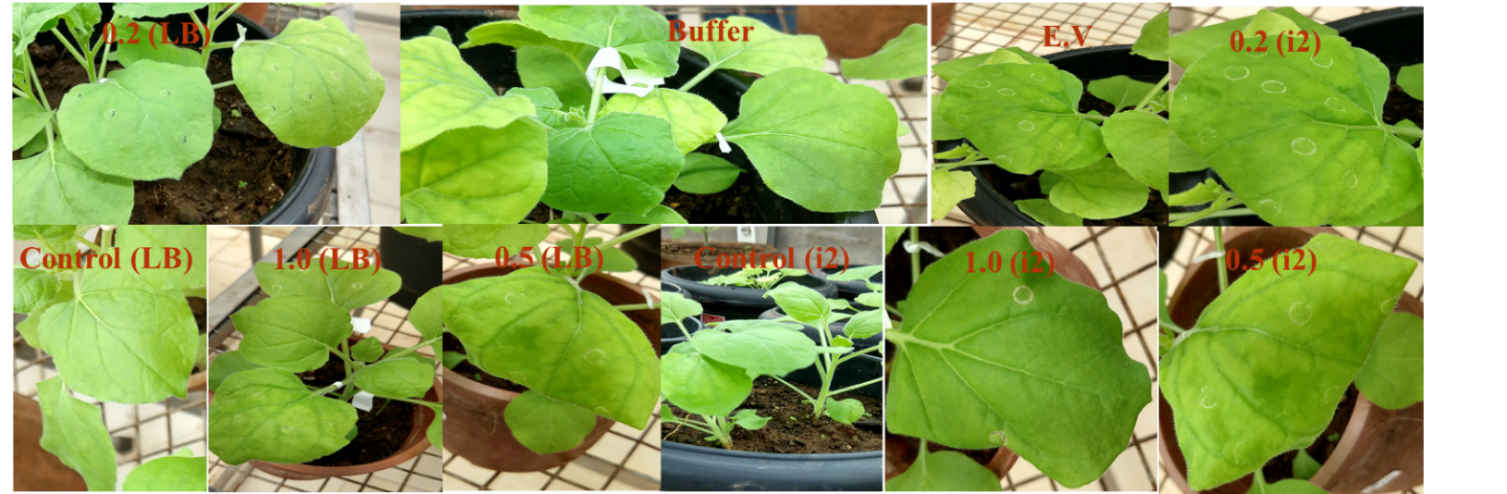

**Fig. S20** Agrobacterium-mediated biotransformation of EutTTS4 (i2) and EutTTS5 (LB) cloned in pRI101 in *Nicotiana Benthamina* (tobacco plant).

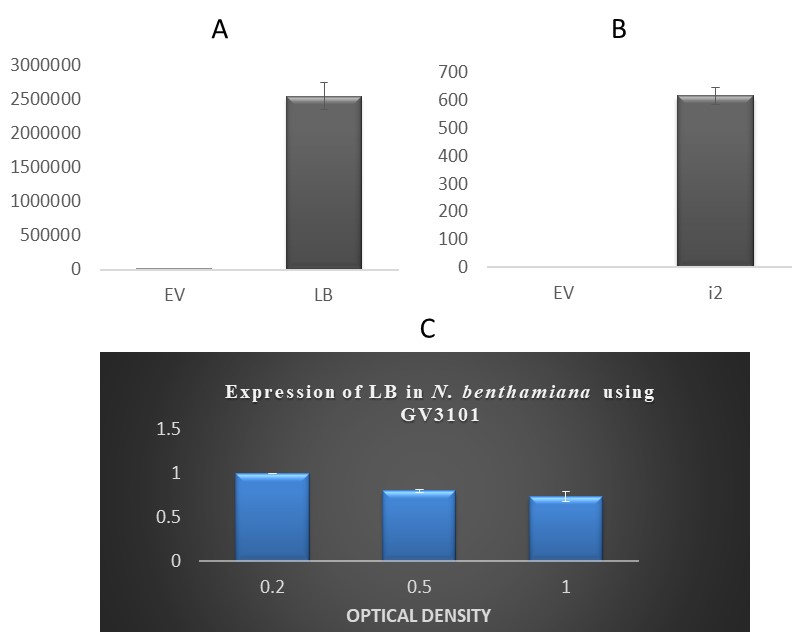

**Fig. S21** Transient transformation expression analysis. (A) Real-time PCR analysis of LB (EugTTS5) using cDNA synthesized from the transiently transformed leaf of *N. benthamiana.* (B) Real-time PCR analysis of i2 (EutTTS4) using cDNA synthesized from the transiently transformed leaf of *N. benthamiana.* (C) Optical density optimization of the agroinfiltrated culture harboring EugTTS5 to obtain maximum overexpression in the leaf of *N. benthamiana.*

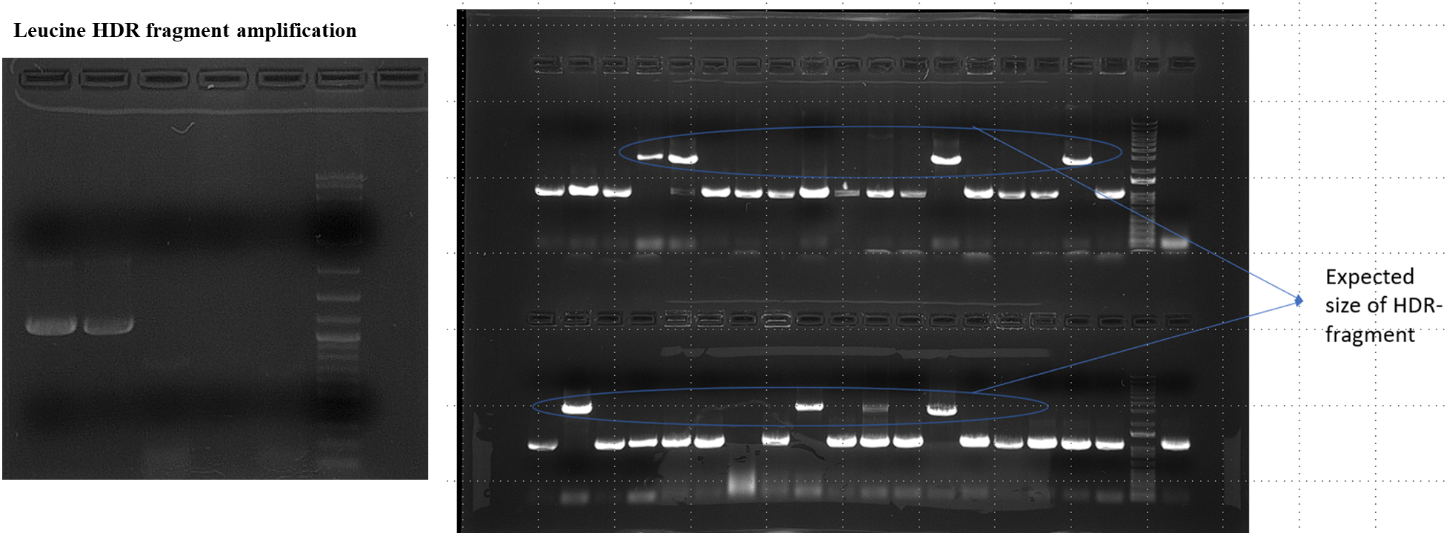

**Fig. S22** Amplification of the LEU2 HDR fragment and the colony PCR of 38 colonies with Seq_hem1 primers.

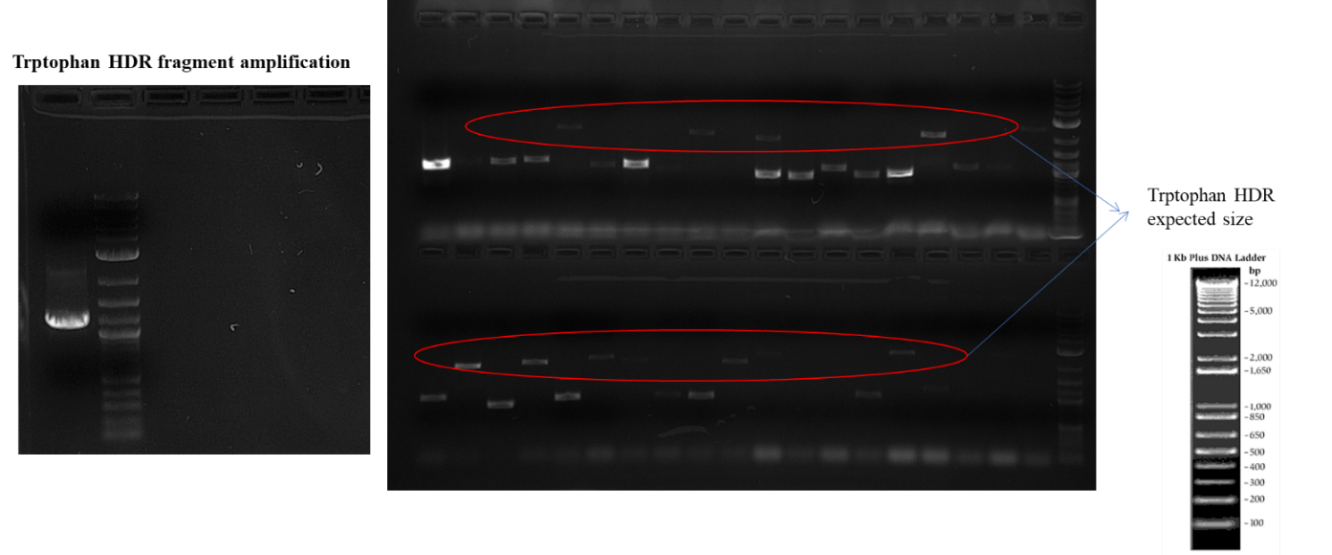

**Fig. S23** Amplification of the Trp1 cassette from the pESC-Trp and the colony PCR of 38 colonies with Seq_hem1 and Seq_erg7 primers.

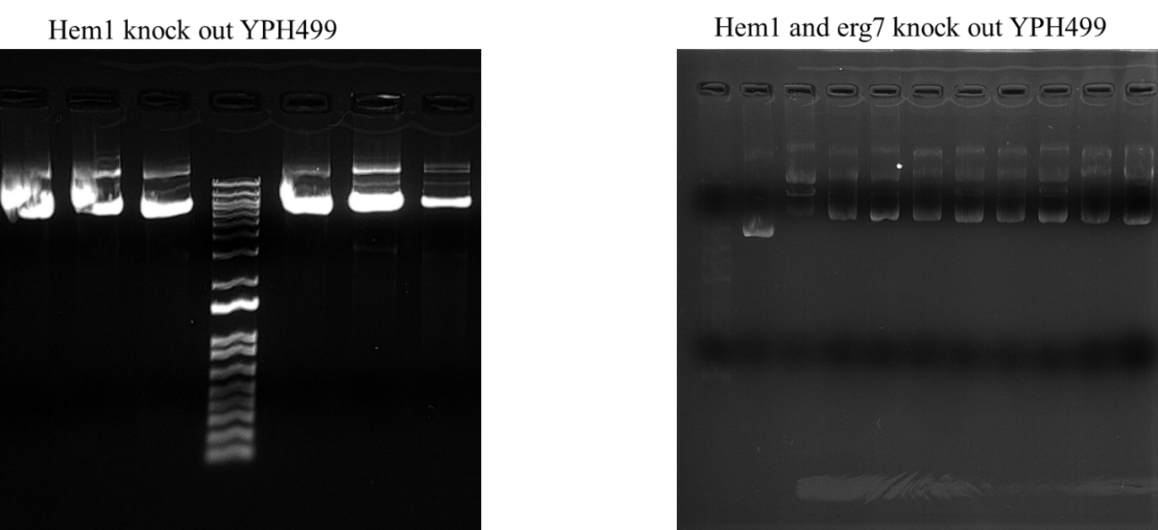

**Fig. S24** Yeast genomic DNA isolated from hem1 mutant and hem1 and erg7 mutant colonies.

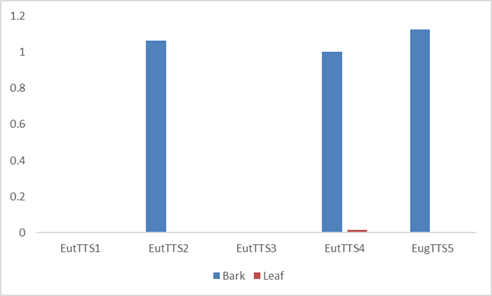

**Fig. S25** Real-time expression analysis of the triterpene synthases .

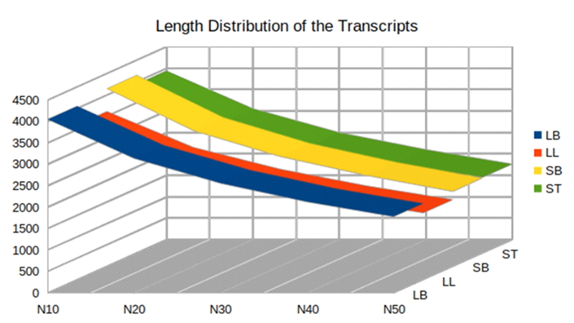

**Fig S26**. Quality check of the assembled transcripts by N10- N50 values.

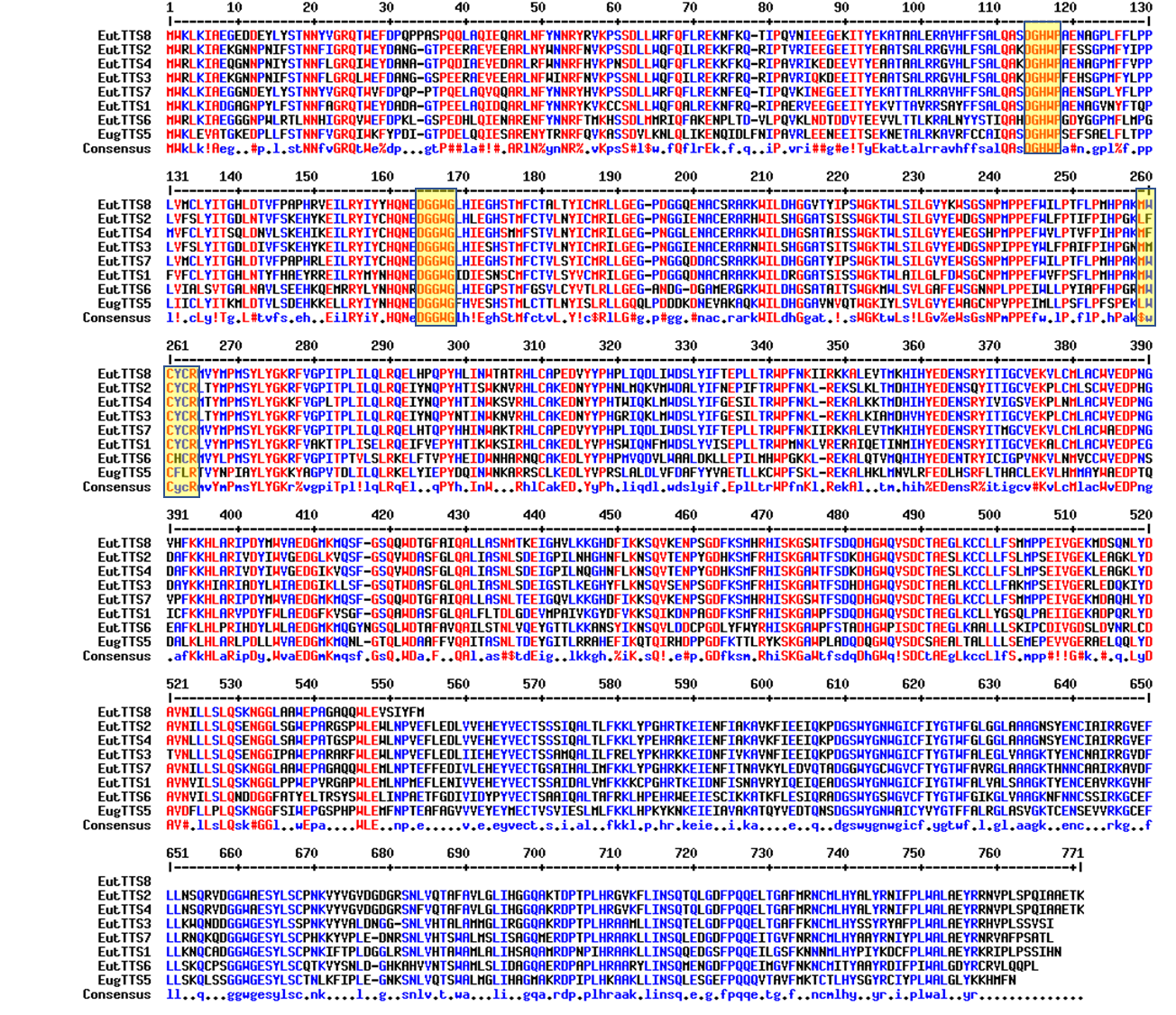

**Fig S27**. Active site amino acid analysis of the triterpene synthases.

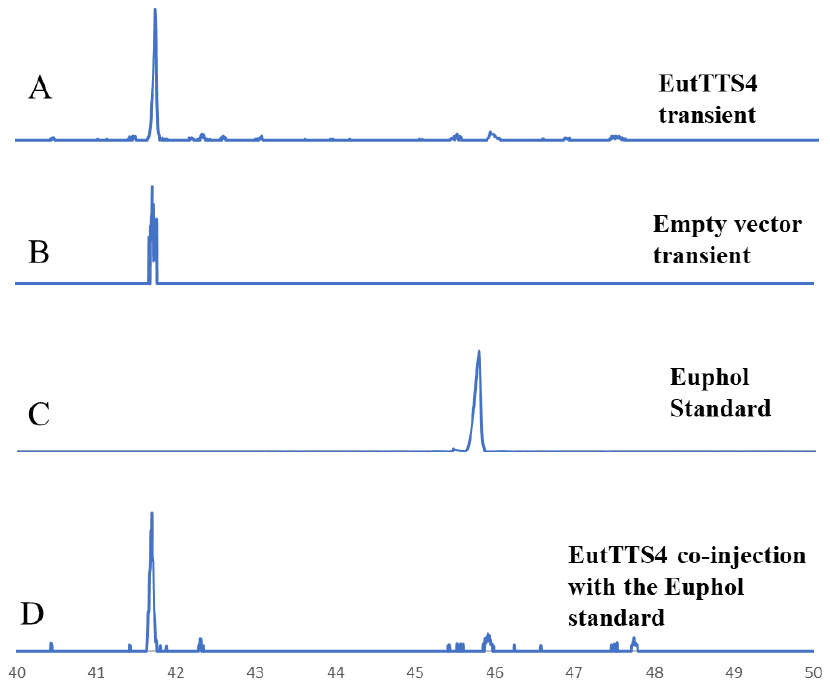

Fig S28. GC-MS chromatogram comparison of the EutTTS4 transient transformation in *N. benthamiana* along with the euphol triterpene standard and co-injection study. A) EIC411 of total metabolite analysis after transient. B) Transient transformation of pRI101-AN empty vector into the *Nicotiana benthamiana*. C) Euphol standard. D) Co-injection study of EutTTS5 product with Euphol standard.

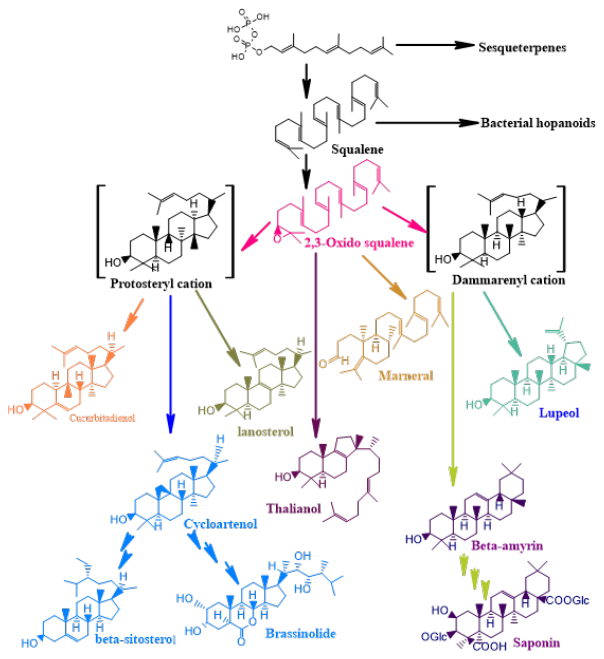

Fig S29. Triterpene biosynthesis is derived from two different cations as intermediate.

| **Transcript ID** | **Description of the transcript** | **Identity** | **Query coverage** | **Transcript length** |
| --- | --- | --- | --- | --- |
| LB_c26609_g1_i1 | beta-amyrin synthase [*Eleutherococcus senticosus*] | 71 | 95 | Miss-assembly  Full length |
| LB_c34999_g1_i1 | beta-amyrin synthase [*Euphorbia tirucalli*] | 96 | 84 | Full length |
| LB_c35711_g3_i1 | cycloartenol synthase [*Ricinus communis*] | 74 | 96 | 900bp from N and 180bp from C |
| LB_c35711_g4_i1 | cycloartenol synthase [*Ricinus communis*] | 85 | 91 | N-terminal 390bp |
| LB_c35995_g1_i3 | beta-amyrin synthase [*Panax quinquefolius*] | 60 | 99 | N-terminal- 1kb |
| LB_c35995_g3_i3 | beta-amyrin synthase [*Bupleurum chinense*] | 57 | 85 | Full length |
| LB_c36778_g2_i1 | putative oxidosqulene cyclase [*Euphorbia tirucalli*] | 88 | 99 | C-terminal 300 bp and N-33bp |
| LB_c36778_g2_i3 | lupeol synthase [*Ricinus communis*] | 77 | 88 | Full length |
| LB_c36778_g2_i10 | lupeol synthase [*Ricinus communis*] | 75 | 88 | Full length |

**Table S1** Triterpene synthases from the *E. grantii* stem tissue.

| **Transcript I.D** | **Description of the transcript** | **Identity** | **Query coverage** | **Transcript length** |
| --- | --- | --- | --- | --- |
| LL_c40919_g7_i5 | beta-amyrin synthase [*Eleutherococcus senticosus*] | 58.8 | 84.58 | Full length |
| LL_c41931_g2_i2 | beta-amyrin synthase [*Euphorbia tirucalli*] | 91.2 | 72.61 | 360 bp from C-terminal |
| LL_c45426_g6_i2 | putative oxidosqulene cyclase [*Euphorbia tirucalli*] | 73.2 | 55.24 | 100 bp is missing from C-terminal |
| LL_c45609_g2_i2 | putative oxidosqulene cyclase [*Euphorbia tirucalli*] | 78.6 | 97.24 | 600 bp missing from C-terminal |

**Table S2** triterpene synthases identified from the *E. grantii* tissue leaf.

| **Transcript I.D** | **Description of the transcript** | **Identity** | **Query coverage** | **Transcript length** |
| --- | --- | --- | --- | --- |
| SB_c34752_g1_i2 | beta-amyrin synthase [*Euphorbia tirucalli*] | 96.1 | 84.29 | full length |
| SB_c35689_g2_i1 | lupeol synthase [*Ricinus communis*] | 85.4 | 99.77 | 600 bp from N-terminal and 450 bp from C-terminal |
| SB_c35742_g1_i1 | cycloartenol synthase [*Ricinus communis*] | 77 | 86.49 | 360 bp from N-terminal and 100 bp from C-terminal |
| SB_c35742_g1_i3 | cycloartenol synthase [*Ricinus communis*] | 85.5 | 95.53 | 100 bp from C-terminal |
| SB_c36358_g2_i4 | beta-amyrin synthase [*Panax quinquefolius*] | 61.7 | 99.55 | 510 bp from N-terminal |
| SB_c36358_g3_i6 | beta-amyrin synthase [*Bupleurum chinense*] | 63.8 | 91.05 | full length |
| SB_c36763_g3_i1 | lupeol synthase [*Ricinus communis*] | 90.5 | 88.79 | full legth |
| SB_c36763_g3_i6 | lupeol synthase [*Ricinus communis*] | 83.6 | 96.05 | 180 bp from N-terminal and 150 bp from C-terminal |
| SB_c36763_g3_i3 | lupeol synthase [*Ricinus communis*] | 83.6 | 96.05 | Full length |

**Table S3** triterpene synthases identified from the *E. tirucalli* stem.

| **Transcript I.D** | **Description of the transcript** | **Identity** | **Query coverage** | **Transcript length** |
| --- | --- | --- | --- | --- |
| ST_c25886_g1_i1 | cycloartenol synthase [*Ricinus communis*] | 86.2 | 82.65 | full length |
| ST_c28604_g2_i2 | beta-amyrin synthase [*Lotus japonicus*] | 82.1 | 95.25 | full length |
| ST_c28940_g2_i2 | beta-amyrin synthase [*Euphorbia tirucalli*] | 95.9 | 86.11 | 600 bp is missing from N-terminal |
| ST_c28338_g1_i1 | beta-amyrin synthase [*Bupleurum chinense*] | 63.8 | 86.67 | full length |
| ST_c27781_g2_i1 | lupeol synthase [*Ricinus communis*] | 97.4 | 83.05 | full length |
| ST_c27781_g2_i2 | lupeol synthase [*Ricinus communis*] | 76.9 | 87.96 | full length |
| ST_c27781_g2_i5 | lupeol synthase [*Ricinus communis*] | 79.3 | 87.63 | full length |
| ST_c27781_g2_i6 | putative oxidosqulene cyclase [*Euphorbia tirucalli*] | 79.4 | 45.72 | full length |
| ST_c14554_g2_i1 | beta-amyrin synthase [*Lotus japonicus*] | 71.2 | 99.72 | 480 bp from N-terminal and 1.3 kb from C -terminal |
| ST_c13055_g1_i1 | putative oxidosqulene cyclase [*Euphorbia tirucalli*] | 80.3 | 100 | 510 bp from N-terminal and 1.2 kb from C- terminal |

**Table S4** triterpene synthases identified from the *E. tirucalli* leaf.

| Real-time primers |  |
| --- | --- |
| LL_c43746_g2_i1 (HMGCR) F | GGTATTGCTGGTCCTTTGTTGCTTAATG |
| LL_c43746_g2_i1 (HMGCR) R | ACAGGTGCTCTTGTCATCCCATCTT |
| ST_c11262_g2_i1 (HMGCR) F | GGGGCTCTTGGTGGATTTAATGCTC |
| ST_c11262_g2_i1 (HMGCR) R | CGGAAGGCATGGTTACAGAAATGTGGAG |
| LL_c43746_g4_i1 (HMGCR) F | GCAATGGGGATGAACATGGTTTCCA |
| LL_c43746_g4_i1 (HMGCR) R | ACTGCCTCGCAAACGACAGATTTAC |
| SB_c32143_g2_i1 (DXS) F | AATGGTTCAGAACTGCCTCAAAGCC |
| SB_c32143_g2_i1 (DXS) R | CGAATCCTCCGATAGAGCCTTCCTC |
| LL_c45552_g1_i2 (DXS) F | AATGGTTCAGAACTGCCTCAAAGCC |
| LL_c45552_g1_i2 (DXS) R | GATTGACCCTTCCTCGACTGTGACT |
| LL_c39969_g3_i2 (DXS) F | GGGCATCACTGTCAGAAAGAGGA |
| LL_c39969_g3_i2 (DXS) R | TTAACTCGACAACTCCAAGGCTCGA |
| LL_c34486_g1_i1 (HMGCR) F | TATTGAGGTGGGTACAGTAGGGGGTG |
| LL_c34486_g1_i1 (HMGCR) R | GGAAGGGGTTCAGGAAGTTAATTTGGACA |
| SB_c32143_g1_i1 (DXS)F | ATGGGCACTGCTGCTGCTCA |
| SB_c32143_g1_i1 (DXS)R | CCCACACCTATACTTGATGTAGTTGAAAACCCTATTC |
| SB_c33639_g2_i1 (DXS) F | TGGAGGGCAGACACATGAAATAGCT |
| SB_c33639_g2_i1 (DXS) R | CTACGACTTTCTCATCCGCTTGTGC |
| LL_c37151_g2_i1 (DXS) F | TGCTTCTGGCTCTACCCTCTTTGAG |
| LL_c37151_g2_i1 (DXS) R | AGTAATTGAGACCTGTCCCACCACC |
| LB_c27998_g1_i1 (DXS) F | ATTGAAGGGGAGAGAGTGGCTTTGT |
| LB_c27998_g1_i1 (DXS) R | GATGGAGTTAGACCTGCCTCAACCA |
| eEF2F | GTGATGTCCGTATGACAGATACCCG |
| eEF2R | CAGACACCCTCAATACAGTCCACC |
| SB_c32143_g1_i1 (DXS) R | GAATAGGGTTTTCAACTACATCAAGTATAGGTGTGGG |
| SB_c33639_g2_i1 (DXS) F | TGGAGGGCAGACACATGAAATAGCT |
| SB_c33639_g2_i1 (DXS) R | CTACGACTTTCTCATCCGCTTGTGC |

**Table S5** primers for the real time PCR analysis of DXS and HMGR identified from *E. grantii* and *E. tirucalli* transcripts*.*

| Primers for cloning |  |
| --- | --- |
| ST_c28604_SacIF | ATGATAGAGCTCAACACAATGTCCATGTGGAAGCTGAAGATAGCA |
| ST_c28604_XbaIR | TGTGAGTCTAGAATTGTGGATGGAAGATGGCAAAGGTATG |
| ST_c27781_i2kpnIF | CTCGGCGGTACCAACACAATGTCCATGTGGAGGCTTAAAATAGCTGAGC |
| ST_c27781_i2XbaIR | AGTGAGTCTAGATTTTGTTTCTGCAGCAATCTGTGGCGATAGAGGAAC |
| SB_c36763_BamHIF | GAAAGGGGATCCAACACAATGTCCATGTGGAGGCTTAAGATAGCAGAG |
| SB_c36763_XbaIR | GCGCGATCTAGAAATACTTACTGAAGACAAAGGAACATGTCGACGA |
| LB_c27781_i6*BamHI*F | ATTCGAGGATCCAACACAATGTCCATGTGGAGGCTCAAAATAGC |
| LB_c27781_i6XbaI | ACGTATTCTAGAAATACTTACTGAAGACAAAGGAACATGTCGAC |
| LB_c35995_g3_KpnI | GATTACGGTACCAACACAATGTCCATGTGGAAGCTAGAGGTAGCT |
| LB_c35995_g3_XhoI | TAGTGTCTCGAGATTAAACATATGTTTTTTGTATAAACCCAGAGC |

**Table S6** Primers for the cloning of EutTTS and EugTTS transcript in pYES2/CT vector.

| TMBL17 generation primers through CRISPR- cas9 | |
| --- | --- |
| ERG7_Trporf_F | atcggtctaccaaagacagatccacgtctttggagactgaGCGGCATCAGAGCAGATTGTACTGA |
| ERG7_Trporf_R | gaaagtggatggtgggtcgtttgcggcttgctgaggggttCTCCTTACGCATCTGTGCGG |
| Seq_hemF | Gctccatttttgcgaggttcggtaactc |
| Seq_hemR | cttcgtaatcgaaacccgactcctggg |
| Seq_erg7F | gacagaattttattctgacacaatcggtctacc |
| Seq_erg7R | TACCTGAGTCAGGCTCTTGAAGCAG |
| Hem_leu orf_F | taactcctctgccgctgtttccacactgaataggctgtccGTGTGGTGCCCTCCTCCT |
| Hem_leu orf_R | gctgtggcagtggcagcagcggcaccagcaccagtagcggCGGTCTAAGGCGCCTGATTCAAG |
| hem1Oligo1 | GATCGCCATTTTTCGCATGTGGTGGTTTTAGAGCTAG |
| hem1Oligo2 | CTAGCTCTAAAACCACCACATGCGAAAAATGGC |
| erg7Oligo1 | GATCTGAGCTAGGCCGAGAAAGCTGTTTTAGAGCTAG |
| erg7Oligo2 | CTAGCTCTAAAACAGCTTTCTCGGCCTAGCTCA |

**Table S7** Primers for the generation of gRNA and sequencing though the genome of TMBL-17.

| Vector sequencing primers | |
| --- | --- |
| CYC_reverse | AGGGCGTGAATGTAAGCGTGAC |
| T7_F primers | TAATACGACTCACTATAGGG |
| pRI101 F | GCACAATCCCACTATCCTTCGC |
| pRI101 R | GCTTCCGGCTCGTATGTTGTG |

**Table S8** Primers for the sequencing of the vector constructs.

| Primers for the expression of EutTTS4 (i2) and EugTTS5 (LB) in *Nicotiana benthamiana* | |
| --- | --- |
| F_EugTTTS5_salI | ATGTAA**GTCGAC**ATGTGGAAGCTAGAGGTAGCTACT |
| R_EugTTS5_kpn1 | GAATTA**GGTACC**ATTAAACATATGTTTTTTGTATAAA |
| F_EutTTS4_kpnI | ATTAAT**GGTACC**ATGTGGAGGCTTAAAATAGCTGA |
| R_EutTTS4_SacI | ATATAT**GAGCTC**TTTTGTTTCTGCAGCAATCTGTGGCGATA |
| ST_c27781_g2_i2_F (EutTTS4) | CAAAGGGGCATGGACATTTTCTGAT |
| ST_c27781_g2_i2_R (EutTTS4) | GCTGATGAAGTGCATTCCACATACTCGT |
| LB_c35995_RTF (EugTTS5) | CATGTAGAGAGCCATAGTACAATGCTATGC |
| LB_c35995_RTR (EugTTS5) | TCCTGCCCACTCGTATACTCC |
| FBOX_nicotianaF | GGCACTCACAAACGTCTATTTC |
| FBOX_nicotianaR | ACCTGGGAGGCATCCTGCTTAT |

**Table S9** Primers for the cloning of EutTTS4 and EugTTS5 transcript in pRI101 vector and for the real-time PCR analysis.

**S1 Method optimized Flow-chart**

**Methods S1. Favarogen genomic DNA isolation steps**

For the yeast genomic DNA isolation, we have used FavorPrep™ Plant Genomic DNA Extraction Mini Kit (Cat no. FAPGK001) In that we have modified the cell lysis as yeast cells have been used.

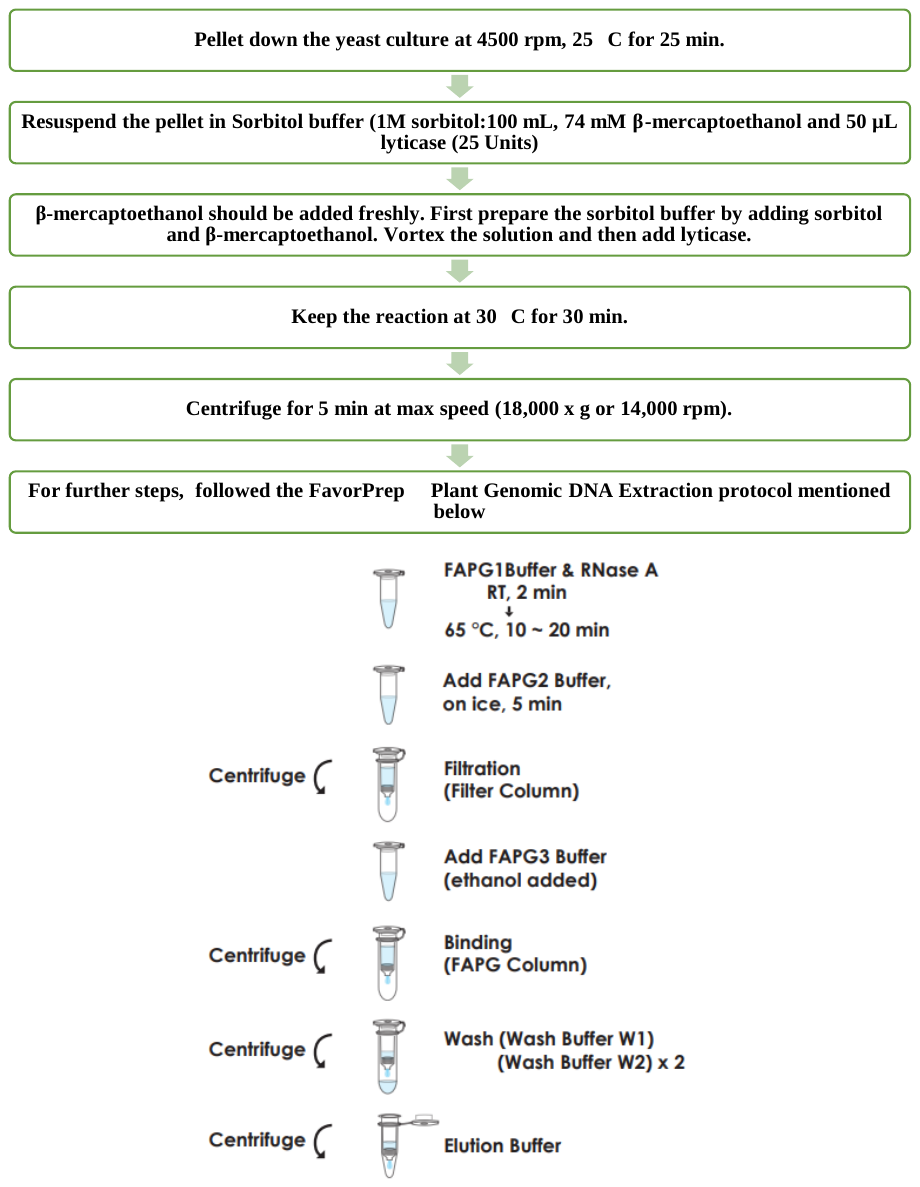

**Methods S2. Yeast Colony PCR steps**

Yeast Colonies were suspended with lyticase (25 units) prepared in PBS, and the whole mixture was kept at 30 oC for 30 min

All PCR components of LA polymerase kit sigma were added to the PCR tube, and 9 µL of PCR mixture was aliquoted

In this PCR mixture, LA polymerase was increased (2.5 µL) for a 50 µL reaction.

1 µL of yeast cell lysis mixture was added in 9 µL of PCR mixture as a template.

PCR was carried using following program: 95oC 5 min, (95oC 30sec, 54oC 30 sec, 68oC)×35, 68oC 10min.

Positive colonies were grown in YPD broth having 13µg/ml hemin chloride and 20µg/ml ergosterol at 30 for 4-5 days

This culture was pelleted at 3000 rpm for 10 minutes, and genomic DNA was isolated using the modified method

>HEM1_position_77 
atgcaacgctccatttttgcgaggttcggtaactcctctgccgctgtttccacactgaataggctgtccacgacagCCGCACCACATGCGAAAAATGGCtatgccaccgctactggtgctggtgccgctgctgccactgccacagcgtcatcaacacatgcagcagcagcagcagccgctgctgccaaccattccacccaggagtcgggtttcgattacgaaggcctgatagattccgaactgcagaagaaaagacttgacaaatcgtacagatatttcaacaatatcaaccgattggccaaggagttccccctagctcatcgccagagagaggcggacaaggtcaccgtttggtgttccaacgactatttagcactttccaagcaccctgaggtattggacgccatgcataaaactatcgacaagtatggttgtggtgccggtggtacaagaaacattgctggccataacatccccactttgaatctggaagccgaattggccactttacacaagaaggaaggtgccttagttttttcgtcatgttacgtagccaacgatgccgtcttatccctactgggtcaaaagatgaaggacttggtgattttctccgacgaactcaaccatgcgtccatgattgtcggtattaagcatgctaacgtaaaaaaacacattttcaaacataatgacttgaacgaattggaacaactgctccagtcataccccaaatccgttcctaaactaattgctttcgaatcagtatattctatggccggttcagtggccgacatagaaaaaatttgcgacttggccgacaaatacggtgctttgaccttcttggatgaagtacatgcggtcggcctgtacggccctcacggtgcaggtgttgcagaacattgtgattttgaaagtcaccgtgcaagtggtattgctaccccaaagaccaatgacaagggcggcgcgaagactgtgatggaccgtgtcgacatgatcaccggcactttaggtaagtctttcggtagcgtaggtggctacgtcgcagcctctaggaaattgatcgattggttcagatcgtttgcacctggtttcattttcaccacgactttaccaccttcagttatggcaggcgctaccgcagcaattagataccaacgttgccacatcgacctaagaacctcgcaacagaaacataccatgtacgtaaagaaagctttccatgagttgggcattccagttattccaaatccttctcatatcgtcccagtgttgattggtaatgctgatttggctaagcaagcttctgacatcttaatcaataagcatcaaatctacgtacaagctatcaacttccctacggttgctcgcggtaccgaaagattgagaattaccccaacgccaggtcacaccaacgatttatctgacatcttaatcaatgcagttgatgatgtgttcaatgagctacagttaccacgtgtcagagactgggaaagccaaggtggcttattgggtgttggagagagcggatttgtggaagagtctaacttatggacatcaagccaactatctttaactaatgacgacttgaaccctaatgttagagaccccatcgttaaacaactagaggtttctagtggtatcaagcagtaa

color regions are integration region for the colony 19 and 21

>ERG7_position_72 
atgacagaattttattctgacacaatcggtctaccaaagacagatccacgtctttggagactgagaactgaTGAGCTAGGCCGAGAAAGCTGGGaatattt**aacccctcagcaagccgcaaa**cgacccaccatccactttcacgcagtggcttcttcaagatcccaaatttcctcaacctcatccagaaagaaataagcattcaccagatttttcagccttcgatgcgtgtcataatggtgcatcttttttcaaactgcttcaagagcctgactcaggtatttttccgtgtcaatataaaggacccatgttcatgacaatcggttacgtagccgtaaactatatcgccggtattgaaattcctgagcatgagagaatagaattaattagatacatcgtcaatacagcacatccggttgatggtggctggggtctacattctgttgacaaatccaccgtgtttggtacagtattgaactatgtaatcttacgtttattgggtctacccaaggaccacccggtttgcgccaaggcaagaagcacattgttaaggttaggcggtgctattggatcccctcactggggaaaaatttggctaagtgcactaaacttgtataaatgggaaggtgtgaaccctgcccctcctgaaacttggttacttccatattcactgcccatgcatccggggagatggtgggttcatactagaggtgtttacattccggtcagttacctgtcattggtcaaattttcttgcccaatgactcctcttcttgaagaactgaggaatgaaatttacactaaaccgtttgacaagattaacttctccaagaacaggaataccgtatgtggagtagacctatattacccccattctactactttgaatattgcgaacagccttgtagtattttacgaaaaatacctaagaaaccggttcatttactctctatccaagaagaaggtttatgatctaatcaaaacggagttacagaatactgattccttgtgtatagcacctgttaaccaggcgttttgcgcacttgtcactcttattgaagaaggggtagactcggaagcgttccagcgtctccaatataggttcaaggatgcattgttccatggtccacagggtatgaccattatgggaacaaatggtgtgcaaacctgggattgtgcgtttgccattcaatactttttcgtcgcaggcctcgcagaaagacctgaattctataacacaattgtctctgcctataaattcttgtgtcatgctcaatttgacaccgagtgcgttccaggtagttatagggataagagaaagggggcttggggcttctcaacaaaaacacagggctatacagtggcagattgcactgcagaagcaattaaagccatcatcatggtgaaaaactctcccgtctttagtgaagtacaccatatgattagcagtgaacgtttatttgaaggcattgatgtgttattgaacctacaaaacatcggatcttttgaatatggttcctttgcaacctatgaaaaaatcaaggccccactagcaatggaaaccttgaatcctgctgaagtttttggtaacataatggtagaatacccatacgtggaatgtactgattcatccgttctggggttgacatattttcacaagtacttcgactataggaaagaggaaatacgtacacgcatcagaatcgccatcgaattcataaaaaaatctcaattaccagatggaagttggtatggaagctggggtatttgttttacatatgccggtatgtttgcattggaggcattacacaccgtgggggagacctatgagaattcctcaacggtaagaaaaggttgcgacttcttggtcagtaaacagatgaaggatggcggttggggggaatcaatgaagtccagtgaattacatagttatgtggatagtgaaaaatcgctagtcgttcaaaccgcatgggcgctaattgcacttcttttcgctgaatatcctaataaagaagtcatcgaccgcggtattgaccttttaaaaaatagacaagaagaatccggggaatggaaatttgaaagtgtagaaggtgttttcaaccactcttgtgcaattgaatacccaagttatcgattcttattccctattaaggcattaggtatgtacagcagggcatatgaaacacatacgctttaa

>LEU2_cassette

GTGTGGTGCCCTCCTCCTTGTCAATATTAATGTTAAAGTGCAATTCTTTTTCCTTATCACGTTGAGCCATTAGTATCAATTTGCTTACCTGTATTCCTTTACTATCCTCCTTTTTCTCCTTCTTGATAAATGTATGTAGATTGCGTATATAGTTTCGTCTACCCTATGAACATATTCCATTTTGTAATTTCGTGTCGTTTCTATTATGAATTTCATTTATAAAGTTTATGTACAAATATCATAAAAAAAGAGAATCTTTTTAAGCAAGGATTTTCTTAACTTCTTCGGCGACAGCATCACCGACTTCGGTGGTACTGTTGGAACCACCTAAATCACCAGTTCTGATACCTGCATCCAAAACCTTTTTAACTGCATCTTCAATGGCCTTACCTTCTTCAGGCAAGTTCAATGACAATTTCAACATCATTGCAGCAGACAAGATAGTGGCGATAGGGTCAACCTTATTCTTTGGCAAATCTGGAGCAGAACCGTGGCATGGTTCGTACAAACCAAATGCGGTGTTCTTGTCTGGCAAAGAGGCCAAGGACGCAGATGGCAACAAACCCAAGGAACCTGGGATAACGGAGGCTTCATCGGAGATGATATCACCAAACATGTTGCTGGTGATTATAATACCATTTAGGTGGGTTGGGTTCTTAACTAGGATCATGGCGGCAGAATCAATCAATTGATGTTGAACCTTCAATGTAGGGAATTCGTTCTTGATGGTTTCCTCCACAGTTTTTCTCCATAATCTTGAAGAGGCCAAAAGATTAGCTTTATCCAAGGACCAAATAGGCAATGGTGGCTCATGTTGTAGGGCCATGAAAGCGGCCATTCTTGTGATTCTTTGCACTTCTGGAACGGTGTATTGTTCACTATCCCAAGCGACACCATCACCATCGTCTTCCTTTCTCTTACCAAAGTAAATACCTCCCACTAATTCTCTGACAACAACGAAGTCAGTACCTTTAGCAAATTGTGGCTTGATTGGAGATAAGTCTAAAAGAGAGTCGGATGCAAAGTTACATGGTCTTAAGTTGGCGTACAATTGAAGTTCTTTACGGATTTTTAGTAAACCTTGTTCAGGTCTAACACTACCGGTACCCCATTTAGGACCAGCCACAGCACCTAACAAAACGGCATCAACCTTCTTGGAGGCTTCCAGCGCCTCATCTGGAAGTGGGACACCTGTAGCATCGATAGCAGCACCACCAATTAAATGATTTTCGAAATCGAACTTGACATTGGAACGAACATCAGAAATAGCTTTAAGAACCTTAATGGCTTCGGCTGTGATTTCTTGACCAACGTGGTCACCTGGCAAAACGACGATCTTCTTAGGGGCAGACATAGGGGCAGACATTAGAATGGTATATCCTTGAAATATATATATATATTGCTGAAATGTAAAAGGTAAGAAAAGTTAGAAAGTAAGACGATTGCTAACCACCTATTGGAAAAAACAATAGGTCCTTAAATAATATTGTCAACTTCAAGTATTGTGATGCAAGCATTTAGTCATGAACGCTTCTCTATTCTATATGAAAAGCCGGTTCCGGCCTCTCACCTTTCCTTTTTCTCCCAATTTTTCAGTTGAAAAAGGTATATGCGTCAGGCGACCTCTGAAATTAACAAAAAATTTCCAGTCATCGAATTTGATTCTGTGCGATAGCGCCCCTGTGTGTTCTCGTTATGTTGAGGAAAAAAATAATGGTTGCTAAGAGATTCGAACTCTTGCATCTTACGATACCTGAGTATTCCCACAGTT AACTGCGGTCAAGATATTTCTTGAATCAGGCGCCTTAGACCG

>erg7trpcassete

AACGACATTACTATATATATAATATAGGAAGCATTTAATAGACAGCATCGTAATATATGTGTACTTTGCAGTTATGACGCCAGATGGCAGTAGTGGAAGATATTCTTTATTGAAAAATAGCTTGTCACCTTACGTACAATCTTGATCCGGAGCTTTTCTTTTTTTGCCGATTAAGAATTAATTCGGTCGAAAAAAGAAAAGGAGAGGGCCAAGAGGGAGGGCATTGGTGACTATTGAGCACGTGAGTATACGTGATTAAGCACACAAAGGCAGCTTGGAGTATGTCTGTTATTAATTTCACAGGTAGTTCTGGTCCATTGGTGAAAGTTTGCGGCTTGCAGAGCACAGAGGCCGCAGAATGTGCTCTAGATTCCGATGCTGACTTGCTGGGTATTATATGTGTGCCCAATAGAAAGAGAACAATTGACCCGGTTATTGCAAGGAAAATTTCAAGTCTTGTAAAAGCATATAAAAATAGTTCAGGCACTCCGAAATACTTGGTTGGCGTGTTTCGTAATCAACCTAAGGAGGATGTTTTGGCTCTGGTCAATGATTACGGCATTGATATCGTCCAACTGCATGGAGATGAGTCGTGGCAAGAATACCAAGAGTTCCTCGGTTTGCCAGTTATTAAAAGACTCGTATTTCCAAAAGACTGCAACATACTACTCAGTGCAGCTTCACAGAAACCTCATTCGTTTATTCCCTTGTTTGATTCAGAAGCAGGTGGGACAGGTGAACTTTTGGATTGGAACTCGATTTCTGACTGGGTTGGAAGGCAAGAGAGCCCCGAAAGCTTACATTTTATGTTAGCTGGTGGACTGACGCCAGAAAATGTTGGTGATGCGCTTAGATTAAATGGCGTTATTGGTGTTGATGTAAGCGGAGGTGTGGAGACAAATGGTGTAAAAGACTCTAACAAAATAGCAAATTTCGTCAAAAATGCTAAGAAATAGGTTATTACTGAGTAGTATTTATTTAAGTATTGTTTGTGCACTTGCCTATGCGGTGTGAAATACCGCACAGATGCGTAAGGAGAAAATACCGCATCAGGAAATTGTAAACGTTAATATTTTGTTAAAATTCGCGTTAAATTTTTGTTAAATCAGCTCATTTTTTAACCAATAGGCCGAAATCGGCAAAATCCCTTATAAATCAAAAGAATAGACCGAGATAGGGTTGAGTGTTGTTCCAGTTTGGAACAAGAGTCCACTATTAAAGAACGTGGACTCCAACGTCAAAGGGCGAAAAACCGTCTATCAGGGCGATGGCCCACTACGTGAACCATCACCCTAATCAAGTTTTTTGGGGTCGAGGTGCCGTAAAGCACTAAATCGGAACCCTAAAGGGAGCCCCCGATTTAGAGCTTGACGGGGAAAGCCGGCGAACGTGGCGAGAAAGGAAGGGAAGAAAGCGAAAGGAGCGGGCGCTAGGGCGCTGGCAAGTGTAGCGGTCACGCTGCGCGTAACCACCACACCCGCCGCGCTTAATGCGCCGCTACAGGGCGCGTCGCGCCATTCGCCATTCAGGCTGCGCAACTGTTGGGAAGGGCGATCGGTGCGGGCCTCTTCGCTATTACGCCAGCTGAATTGGAGCGACCTCATGCTATACCTGAGAAAGCAACCTGACCTACAGGAAAGAGTTACTCAAGAATAAGAATTTTCGTTTTAAAACCTAAGAGTCACTTTAAAATTTGTATACACTTATTTTTTTTATAACTTATTTAATAATAAAAATCATAAATCATAAGAAATTCGCTTATTTAGAAGTGTCAACAACGTATCTACCAACGATTTGACCCTTTTCCATCTTTTCGTAAATTTCTGGCAAGGTAGACAAGCCGACAACCTTGATTGGAGACTTGACCAAACCTCTGGCGAAGAATTGTTAATTAAGAGCTCAGATCTTATCGTCGTCATCCTTGTAATCCATCGATACTAGTGCGGCCGCCCTTTAGTGAGGGTTGAATTCGAATTTTCAAAAATTCTTACTTTTTTTTTGGATGGACGCAAAGAAGTTTAATAATCATATTACATGGCATTACCACCATATACATATCCATATACATATCCATATCTAATCTTACTTATATGTTGTGGAAATGTAAAGAGCCCCATTATCTTAGCCTAAAAAAACCTTCTCTTTGGAACTTTCAGTAATACGCTTAACTGCTCATTGCTATATTGAAGGTTATTACTGAGTAGTATTTATTTAAGTATTGTTTGTGCACTTGCCTATGCGGTGTGAAATACCGCACAGATGCGTAAGGAG
